# ALBA proteins facilitate cytoplasmic YTHDF-mediated reading of m^6^A in plants

**DOI:** 10.1101/2024.06.06.597704

**Authors:** Marlene Reichel, Mathias Due Tankmar, Sarah Rennie, Laura Arribas-Hernández, Martin Lewinski, Tino Köster, Naiqi Wang, Anthony A. Millar, Dorothee Staiger, Peter Brodersen

**Affiliations:** University of Copenhagen, Copenhagen Plant Science Center, Department of Biology, Copenhagen N, Denmark; Department of RNA Biology and Molecular Physiology, Faculty of Biology, Bielefeld University, D-33615 Bielefeld, Germany; Department of Biology, Copenhagen University, Copenhagen N, Denmark; Division of Plant Science, Research School of Biology, The Australian National University, Canberra ACT 2601, Australia

**Author notes:** These authors contributed equally to this work (Marlene Reichel, Mathias Due Tankmar, Sarah Rennie).

**Keywords:** *N6*-methyladenosine (m^6^A), YTHDF proteins, ECT2, ALBA proteins, Intrinsically disordered regions (IDR), Arabidopsis, iCLIP, HyperTRIBE

## Abstract

*N6*-methyladenosine (m^6^A) exerts many of its regulatory effects on eukaryotic mRNAs by recruiting cytoplasmic YT521-B homology domain family (YTHDF) proteins. Here, we show that in Arabidopsis, the interaction between m^6^A and the major YTHDF protein ECT2 also involves the mRNA-binding ALBA protein family. ALBA and YTHDF proteins physically associate via a deeply conserved short linear motif in the intrinsically disordered region of YTHDF proteins, their mRNA target sets overlap, and ALBA4 binding sites are juxtaposed to m^6^A sites. These binding sites correspond to pyrimidine-rich elements previously found to be important for m^6^A binding of ECT2. Accordingly, both biological functions of ECT2 and its binding to m^6^A targets *in vivo* require ALBA association. Our results introduce the YTHDF-ALBA complex as the functional cytoplasmic m^6^A-reader in plants and define a molecular foundation for the concept of facilitated m^6^A reading that increases the potential for combinatorial control of biological m^6^A effects.

## INTRODUCTION

*N6*-methyladenosine (m^6^A) occurs widely in eukaryotic mRNAs. It is introduced into pre-mRNA during transcription in adenosines in DR**A**CH/GG**A**U (D=A/G/U, R=A/G, H=A/C/U) motifs by a deeply conserved RNA polymerase II-coupled methyltransferase complex^1^. m^6^A is required to complete embryogenesis in vertebrates and plants^2,3^. It is also important for yeast sporulation^4^ and for sex determination and neuronal development in insects^5,6^. Many developmental functions of m^6^A rely on cytoplasmic RNA-binding proteins (RBPs) specialized for m^6^A recognition, or “reading”, via a YTH domain^7–10^. These YTH <u>d</u>omain family (YTHDF) m^6^A readers contain the YTH domain at the C-terminus, preceded by a long intrinsically disordered region (IDR)^11^.

Higher plants encode an expanded family of YTHDF proteins with, for instance, 11 members in *Arabidopsis thaliana* (Arabidopsis)^12,13^. They are called EVOLUTIONARILY CONSERVED C-TERMINAL REGION 1-11 (ECT1-11) with reference to the deeply conserved YTH domain at the C-terminus^14^, following an intrinsically disordered region (IDR) of more variable length and sequence. ECT2 and ECT3 are crucial for post-embryonic development^7,15^, as they stimulate cell division in primordial cells^15^. Thus, double knockout of *ECT2* and *ECT3* causes slow formation and aberrant morphology of leaves, roots, stems, flowers and fruits, and these phenotypes are generally exacerbated by additional knockout of *ECT4*^15^. The developmental role of the m^6^A-ECT module is conserved in plants, because knockout of tomato and rice *ECT* genes also causes delayed development^16,17^.

Three features of the molecular functions of ECT proteins that promote growth during organogenesis have been defined. First, they are deeply conserved, because the sole YTHDF protein encoded by the liverwort *Marchantia polymorpha* that diverged from higher plants ∼450 million years ago^18,19^ can functionally replace Arabidopsis ECT2 when expressed in primordial cells in *ect2 ect3 ect4* mutants^20^. Second, ECTs interact with the major cytoplasmic poly(A)-binding proteins PAB2/4/8^21,22^. This interaction is mediated by a conserved tyrosine-rich motif in the IDR of ECT2 and is required for developmental functions of ECT2^21^. Third, most Arabidopsis *ECT* paralogues across phylogenetic subclades retain the ability to complement *ect2 ect3 ect4* mutants upon ectopic expression in primordial cells^20^. For the three Arabidopsis ECT proteins unable to perform this basal function, the divergence can at least in part be ascribed to differences in their N-terminal IDRs^20^, including the loss of the PAB2/4/8-interacting motif^21^. Thus, the molecular properties of the IDRs of ECT proteins are central to understand their biological functions.

At least three distinct molecular properties of IDRs in RBPs are expected to contribute to their functions. First, IDRs often mediate self-assembly such that above a critical concentration, they separate into a phase distinct from the aqueous solution^23^. This is also the case for plant ECT proteins^7,13^, and negative feedback regulation of important stress-related m^6^A-containing mRNAs may indeed rely on ECT-mediated phase separation^24,25^. Second, the IDR may influence RNA-binding activity, either by stabilization of the RNA-bound conformation of the globular RNA-binding domain^26^, or through direct RNA binding activity, as in the case of Arg-Gly-Gly (RGG) repeats in IDRs^27^. The non-RGG-containing IDR of ECT2 may have such properties, because crosslinks to target mRNAs specific to the IDR were identified in crosslinking-immunoprecipitation-sequencing (CLIP-seq) data^28^, and because deletion of the IDR from ECT2 strongly reduces RNA binding capacity *in vivo*^21^. Third, short linear motifs (SLiMs) may be used to mediate direct binding to other proteins^29^, including other RBPs and regulators of the rate of translation and mRNA decay, as in the example of the ECT2-PAB2/4/8 interaction^21^.

The ALBA (<u>a</u>cetylation lowers <u>b</u>inding <u>a</u>ffinity) family of proteins was found in mRNA interactome capture screens to be a prominent group of mRNA-associated RBPs in Arabidopsis^30,31^. The ALBA superfamily of proteins contains an archaeal and two eukaryotic families. Proteins in the Sac10b archaeal family^32^ exhibit acetylation-sensitive DNA-binding activity and have histone-like properties^33–36^, but may also have RNA chaperone functions^37^. The two eukaryotic families group around two distinct subunits of RNaseP/MRP complexes, Rpp20 or Rpp25^32^. Plants encode ALBA proteins belonging to both eukaryotic families. The Rpp20-related forms are short and contain only the ∼95 amino acid globular ALBA domain, while the Rpp25-related forms are long and contain ∼200-300 amino acid C-terminal extensions, often IDRs with many RGG repeats^38^. The sequence similarity within the eukaryotic families is limited, and in most cases, it is not clear whether the ALBA proteins are mRNA-binding or have other RNA-related functions. mRNA-binding ALBA proteins have been studied in the parasitic protist *Trypanosoma brucei* where short and long forms are required for translational regulation of many mRNAs during the transition between mammalian and insect hosts, in particular for growth after commitment to differentiation into the insect-specific form^39,40^.

A requirement of ALBA proteins for growth is recurrent in several plant species^41,42^, first observed in the liverwort *M. polymorpha* where the sole long RGG-repeat-containing ALBA protein is necessary for the development of root-like structures called rhizoids^41^. Arabidopsis encodes three short ALBA proteins in the Rpp20 group, ALBA1-3, and three long ALBA proteins in the Rpp25 group, ALBA4-6^38^. Single knockouts of *ALBA1* and *ALBA2* cause defective root hair development, but no overall growth defects^41^. In contrast, combined knockout of *ALBA4-6* leads to slow seedling development, including defective root growth^43^. A similar defect in root growth was also observed in cotton upon RNAi-mediated knockdown of *ALBA* genes^42^, further supporting the idea that ALBA proteins stimulate tissue growth in plants. Nonetheless, the molecular basis for their growth-promoting function has not been defined.

In this study, we show that ALBAs and ECT2 associate via a deeply conserved SLiM in the IDR of ECT2 to form an efficient m^6^A reader complex in Arabidopsis. The mRNA target sets of ALBA proteins overlap significantly with those of m^6^A-ECT2/3, and ALBA4 binding sites in 3’-UTRs are juxtaposed to m^6^A sites. Finally, ALBA proteins facilitate the association of ECT2 with m^6^A-modified transcripts and are necessary for biological functions of m^6^A-ECT2/3. Thus, our results uncover a mechanism for facilitated m^6^A reading by YTHDF-interacting RBPs with binding sites in close proximity to m^6^A.

## RESULTS

### The N8 IDR element of ECT2 is required for normal growth of leaf primordia

We previously showed that a 37-amino acid residue region in the N-terminal IDR of ECT2, N8, is required for full activity in promoting growth of leaf primordia^21^. Since deletion of the N8-encoding region from an *ECT2-mCherry* gDNA transgene caused a decrease, not abolishment, of the complementation frequency of the *ect2-1 ect3-1 ect4-2* (henceforth, *te234*) triple knockout mutant^7,21^, we first sought to corroborate the importance of N8 by independent means. To this end, we used CRISPR-Cas9 in the *ect3-1 ect4-2* genetic background to generate a chromosomal in-frame *ECT2* deletion matching almost exactly ΔN8 (*ect2-5*, **Figure 1A**, **Figure S1**). The resulting *ect2-5 ect3-1 ect4-2* mutant exhibited slow emergence of the first true leaves, albeit less pronounced than *te234* (**Figure 1B,C**). These results verify that deletion of N8 causes partial loss of ECT2 function. We also confirmed that the ECT2-5 protein accumulated to levels similar to the wild type protein (**Figure 1D**), excluding the possibility that the partial loss of ECT2 function in *ect2-5* mutants is due to decreased dosage.

**Figure 1.**
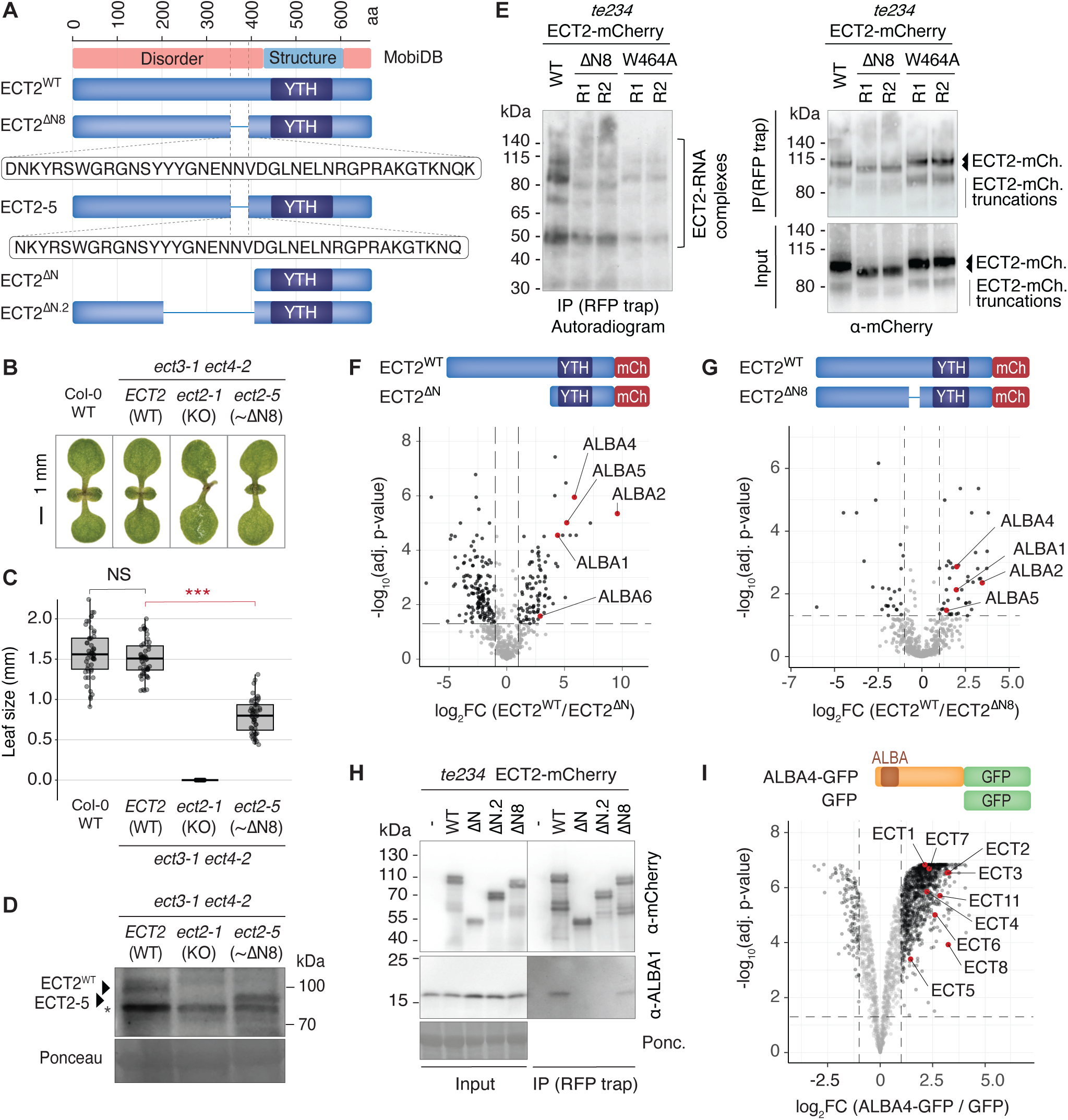
The N8 IDR-element of ECT2 is required for growth promotion, RNA association and interaction with ALBA proteins. **(A)** Schematic representation of wild type and mutant ECT2 proteins. The MobiDB^66^ track (top) displays regions predicted to be structured or disordered. **(B)** Images of representative seedlings of the indicated genotypes taken at 7 days after germination (DAG). **(C)** Quantification of first true leaf size in seedlings of the indicated genotypes 7 DAG. 50 seedlings were measured for each genotype (n = 50). The boxes show the interquartile range (25th–75th percentile), with the central line marking the median. Whiskers extend 1.5 times the interquartile range. Asterisks indicate significance according to *p*-value of *t*-tests between the indicated genotypes. NS, not significant (*** p < 10^-5^). **(D)** Protein blots of total lysates prepared from 12-day old seedlings of the indicated genotypes, probed with ECT2-specific antisera^7^. Arrows indicate the positions of the ECT2^WT^ protein and the ECT2-5 protein containing the N8-like deletion. The asterisk indicates an unspecific band. Ponceau staining serves as the loading control. **(E)** Results of an *in vivo* UV crosslinking ECT2-mCherry-immunoprecipitation experiment, followed by PNK-labelling of precipitated RNA with γ-^32^P-ATP. Left panel, autoradiogram of ^32^P-radiolabelled RNA-protein complexes purified from plants expressing ECT2^WT^-mCherry, ECT2^ΔN^^8^-mCherry or the aromatic cage mutant ECT2^W464A^-mCherry. Molecular weight marker positions and the location of the verified ECT2-mCherry-RNA complexes^28^ are indicated. The presence of several bands of unequal intensity is due to partial proteolysis of the ECT2 IDR during immunoprecipitation and differential labelling efficiency of the different RNPs^28^. Right panels, mCherry immunoblots of the immunoprecipitated (top) and total fractions (input, bottom). Samples were pools of 3 independent lines for each genotype. **(F-G)** Volcano plots showing the differential abundance of proteins co-purified with ECT2-mCherry variants (RFP-trap) measured by mass spectrometry of immunopurified fractions (IP-MS). All ECT2-mCherry variants were expressed in the *te234* mutant background. Diagrams above each plot indicate the proteins compared. Statistical significance was determined using empirical Bayes statistics with Benjamin–Hochberg adjusted P-values. The data underlying the plot in (F) have previously been published^21^. **(H)** Co-immunoprecipitation assay using mCherry immunoprecipitation from 10-day old seedlings expressing the indicated ECT2-mCherry variants (see (A)), followed by immunoblot analysis with mCherry- and ALBA1-specific antibodies. Seedlings from three independent transgenic lines were pooled in this experiment. **(I)** Volcano plots showing the differential abundance of proteins co-purified with ALBA4-GFP as determined by IP-MS from total lysates prepared from 7-day-old seedlings. Statistical significance was calculated using empirical Bayes statistics with Benjamini– Hochberg adjusted p-values.

### N8 is necessary for full RNA association of ECT2

We next conducted *in vivo* UV crosslinking and immunoprecipitation (CLIP) experiments to test whether RNA association was affected by deletion of N8. We quantified crosslinked RNA immunoprecipitated with ECT2^WT^-mCherry or ECT2^ΔN8^-mCherry by polynucleotide kinase (PNK)-mediated radiolabeling, using the previously described assay conditions that allow assignment of the radiolabeled species as ECT2-mCherry-RNA complexes with different sizes resulting from cleavage of the IDR in the lysis buffer^28^. These experiments revealed a reproducible reduction in RNA association of ECT2^ΔN8^-mCherry compared to ECT2-mCherry, albeit less pronounced than the reduction obtained with the m^6^A-binding deficient ECT2^W464A^-mCherry mutant ^28^ (**Figure 1E**). These results suggest that N8 is involved in RNA association, either directly or through interaction with other RBPs whose presence may enhance the affinity of ECT2 for m^6^A-containing mRNAs.

### N8 is necessary for interaction with ALBA proteins

To test whether N8 is required for association of ECT2 with other RBPs, we used comparative immunoprecipitation-mass spectrometry (IP-MS) with stable transgenic lines expressing comparable amounts of either ECT2^WT^-mCherry or ECT2^ΔN8^-mCherry in the *te234* background ([21], **Figure S2A**). We also included three lines of ECT2^ΔN^-mCherry lacking the entire N-terminal IDR ([21], **Figure 1A**) as an additional negative control. All immunopurifications were done in the presence of RNaseA to recover RNA-independent interactors. These experiments revealed that the family of ALBA proteins, in particular ALBA1/2/4/5, were prominent interactors of ECT2 (**Figure 1F**), and that the interaction was strongly dependent on N8 (**Figure 1G, Table S1**).

We used three different approaches to verify the ALBA-ECT interaction and its dependence on N8. First, we raised an antibody specific for ALBA1 (**Figure S2B**) and used it to confirm that ALBA1 enrichment is reduced, but not abolished, upon deletion of N8 (**Figure 1H**). We also included two larger IDR deletion mutants in this experiment, ECT2^ΔN^-mCherry and ECT2^ΔN.2^-mCherry lacking the ∼200 amino acid residues proximal to the YTH domain ([21], **Figure 1A,H**). ALBA1 levels were not detectable in immunopurified fractions of these two mutants (**Figure 1H**), perhaps suggesting that additional determinants of ALBA interaction are located in the IDR outside of the N8 region. Second, inspection of IP-MS data with HA-ECT2 and with tagged versions of the two YTHDF paralogs ECT3 (ECT3-Venus) and ECT1 (ECT1-TFP)^21^, both of which have m^6^A-binding capacity^7,20,24,28,44^, revealed enrichment of ALBA proteins over the negative controls (**Figure S2C**). Third, comparative IP-MS analysis carried out with ALBA4-GFP and free GFP revealed a clear enrichment of several ECT proteins, including ECT1-8 and ECT11, in the ALBA4-GFP purified fractions (**Figure 1I, Figure S2D, Table S1**). These results indicate that ALBA and ECT proteins physically associate *in vivo* and that the ECT2-ALBA association involves the N8 region of the ECT2 IDR. We also take particular note of the combination of two properties. First, deletion of N8 causes reduced RNA binding of ECT2 *in vivo*. Second, ECT interactors of ALBAs include ECT1 and ECT11 which have m^6^A-binding capacity but not the function of ECT2 required for leaf formation^20^. Hence, our results suggest that the ALBA-ECT interaction mediates a molecular property common to all ECT proteins, perhaps m^6^A-binding.

### AlphaFold3 modeling highlights a conserved SLiM in N8 as key for interaction of ECT2 with ALBA domains and RNA

Because many proteins in addition to ALBA1/2/4/5 lose enrichment in immunopurified ECT2 fractions upon deletion of N8 (**Figure 1G**), we sought to further narrow the region in the IDR of ECT2 required for ALBA interaction. We noticed that a SLiM within N8 is conserved both in Arabidopsis ECT paralogs and in ECT2 orthologs from representatives of major clades representing land plant evolution, including *M. polymorpha* YTHDF (**Figure 2A**). Since the N8 region is required for full association of ECT2 with both mRNA and ALBA proteins *in vivo*, we hypothesized that the N8 element might mediate interaction between the three molecules, perhaps via the conserved SLiM. Thus, we used AlphaFold3^45^ to query whether a complex composed of an ALBA-domain dimer^36^, an ECT2 fragment spanning the YTH-domain plus the SLiM-containing proximal part of the IDR, and an m^6^A-containing 10-nt RNA could be modeled. Interestingly, AlphaFold3 generated a model of high confidence overall (**Figure 2B-D**, **Figure S3A-B**). The model features several interactions between the N8-SLiM and the YTH domain, and situates the SLiM centrally between the YTH domain, the ALBA domains, and the m^6^A-containing RNA (**Figure 2B, Figure S3A**). Because these properties offer straight-forward explanations for the reduced ALBA- and RNA-association of ECT2^ΔN8^ *in vivo*, we devoted further efforts to the study of the SLiM and refer to it as the <u>Y</u>THDF-<u>A</u>LBA Interaction <u>M</u>otif (YAIM) in the remainder of this report.

**Figure 2.**
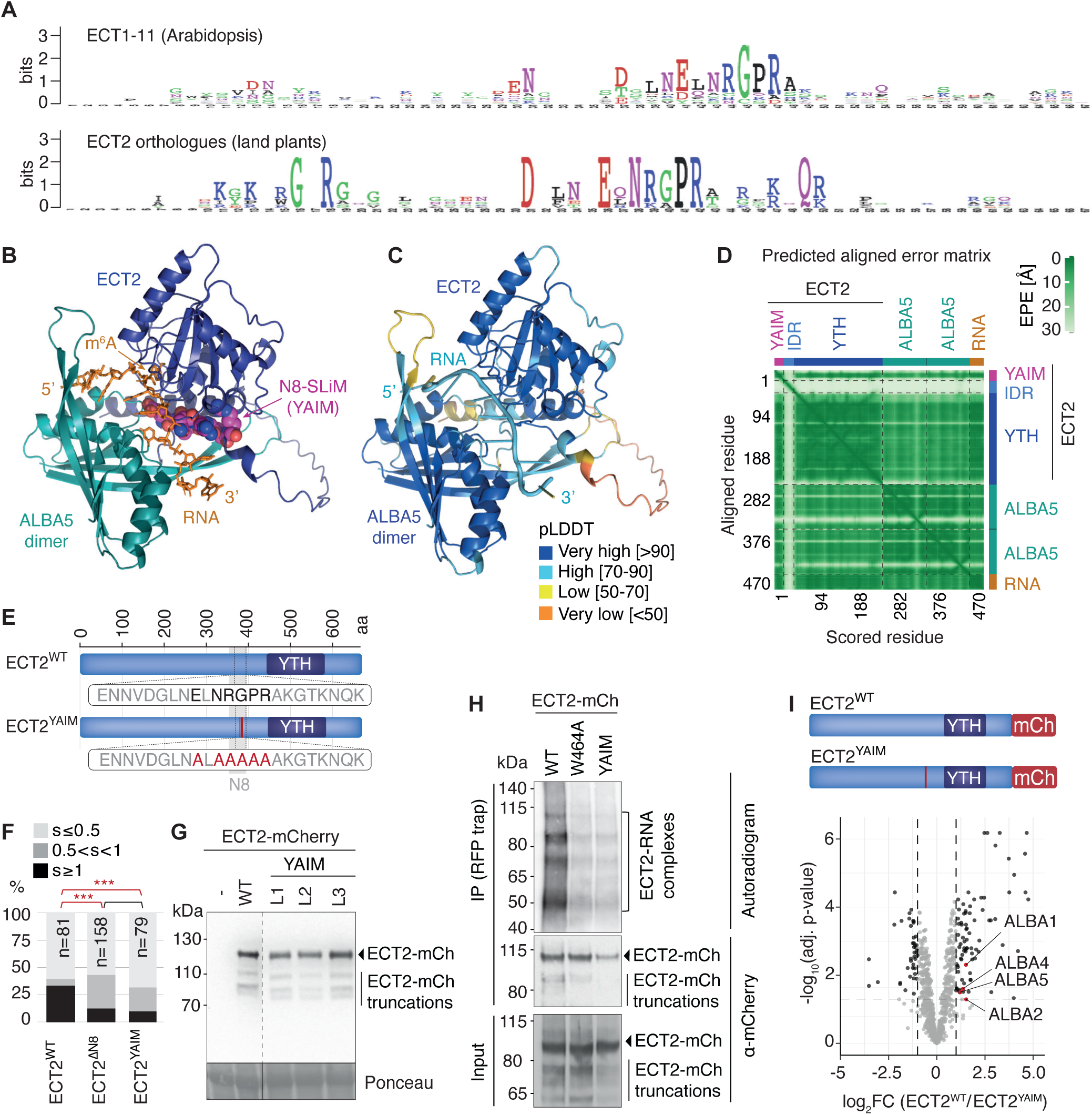
The ECT2-ALBA interaction is mediated by a conserved short linear motif in the N8 element of the ECT2 IDR. **(A)** Logo representations of sequence conservation in the N8 region of the IDR of plant YTHDF proteins. Top, Arabidopsis ECT paralogues (ECT1-ECT11). Bottom, Arabidopsis ECT2 orthologues from 7 different species representing major clades of land plants separated by ∼500 million years of evolution [liver worts (*Marchantia polymorpha*), mosses (*Physcomitrella patens*), lycophytes (*Selaginella moellendorfii*), ferns (*Ceratopteris richardtii*), Amborella (*Amborella trichopoda*), monocots (*Oryza sativa*), dicots (*Arabidopsis thaliana*). Logos^67^ were generated using the Weblogo tool^68^, and sequences were aligned with ClustalW^69^. **(B)** AlphaFold3 model of the complex between ECT2 (YTH domain plus a YAIM-containing fragment of the N-terminal IDR), two ALBA5 subunits (ALBA domains only), and a 10-nt RNA [5’-AAA(m^6^A)CUUCUG-3’]. The YAIM is accentuated in space fill mode (magenta, C; blue, N; red, O), all other protein elements in cartoon mode, and the RNA in stick mode. **(C)** Same view of the model as in panel (B) but colored according to the predicted Local Distance Difference Test (pLDDT) score calculated by Alphafold3 to indicate model confidence on a local per-residue basis^45^. **(D)** 2D plot generated by AlphaFold3 showing the Predicted Aligned Error (PAE) indicating the Expected Position Error (EPE) in Ångströms (white-green scale) in the relative positions of each pair of residues in the complex^45^. The location of subunits and structural elements along the axes is indicated. An additional view of the complex is provided in Supplemental Figure S4. **(E)** Schematic representation of the ECT2^YAIM^ mutant with alanine substitutions in the YTH-ALBA Interaction Motif (YAIM) highlighted in red. **(F)** Categorized leaf size distribution of 9-day-old primary transformants of *te234* mutants expressing wild type or mutant versions of ECT2-mCherry as indicated. Red lines with asterisks denote significant differences based on pairwise Fisher exact tests with Holm-adjusted p-values (*p < 0.05, **p < 0.01, ***p < 0.001). Black line indicates no significant difference. **(G)** Anti-mCherry immunoblot from total lysates of 9-day-old seedlings of transgenic lines expressing either a fully complementing ECT2^WT^-mCherry transgene^7^ or the ECT2^YAIM^-mCherry construct (L1-L3, three independent lines), or without any ECT2 transgene (–), all in the *te234* mutant background. Dashed lines indicate that lanes have been removed for presentation purposes. Ponceau staining is used as a loading control. **(H)** Results of an *in vivo* UV crosslinking ECT2-mCherry-immunoprecipitation experiment, followed by PNK-labelling of precipitated RNA with γ-^32^P-ATP. Top panel, autoradiogram of ^32^P-radiolabelled RNA-protein complexes purified from plants expressing ECT2^WT^-mCherry, the aromatic cage mutant ECT2^W464A^-mCherry, or ECT2^YAIM^-mCherry. Molecular weight marker positions and the location of the verified ECT2-mCherry-RNA complexes^28^ are indicated. The presence of several bands of unequal intensity is due to partial proteolysis of the ECT2 IDR during immunoprecipitation and differential labelling efficiency of the different RNPs^28^. Middle and bottom panels, immunoblots against mCherry showing the ECT2-mCherry proteins in the IP (middle) and total lysates (input, bottom). Samples were pools of 3 independent lines for each genotype. **(I)** Volcano plot showing differential abundance of proteins detected by mass spectrometry in mCherry immunoprecipitates from *te234* seedlings expressing either ECT2^YAIM^-mCherry or ECT2^WT^-mCherry. Statistical significance was determined using empirical Bayes statistics with Benjamini–Hochberg adjusted p-values.

### The YAIM is required for ECT2-ALBA interaction and ECT2 function

We next generated a YAIM mutant of ECT2 containing several alanine substitutions (**Figure 2E**). The ECT2^YAIM^-mCherry mutant exhibited a reduced *te234* complementation frequency similar to ECT2^ΔN8^-mCherry (**Figure 2F**), despite the fact that protein levels in several independent transgenic lines were similar to those obtained with an *ECT2^WT^-mCherry* transgene (**Figure 2G**). These observations demonstrate the *in vivo* importance of the YAIM for ECT2 function. At the molecular level, the ECT2^YAIM^-mCherry mutant also exhibited defects closely resembling those of ECT2^ΔN8^-mCherry: less RNA could be crosslinked and immunoprecipitated with ECT2^YAIM^-mCherry than with ECT2^WT^-mCherry (**Figure 2H**), and ALBA1/2/4/5 were depleted in ECT2^YAIM^-mCherry immunopurifications relative to ECT2^WT^-mCherry (**Figure 2I, Figure S3C, Table S1**). We also used the ALBA1 antibody to verify reduced association with ECT2^YAIM^-mCherry compared to ECT2^WT^-mCherry (**Figure S3D**). Taken together, we conclude that the YAIM is required for ALBA association and for full target RNA-binding of ECT2 *in vivo*, as predicted by the AlphaFold3 model of the (ALBA4)_2_-ECT2-RNA complex. We note, however, that ALBA1/2/4/5 were not specifically depleted from ECT2-mCherry purifications upon mutation of the YAIM, perhaps suggesting that the primary function of the YAIM is to mediate ALBA- and RNA-interaction, and that the ECT2-ALBA-RNA complex generates a platform required for interaction with multiple other proteins.

### A model for concerted m^6^A-ECT-ALBA function in vivo

The results presented so far suggest that ECTs and ALBAs act in concert to bind to m^6^A-sites in mRNA targets. A basic prediction of this hypothesis is that ECT2 and ALBAs are expressed in the same cells. Examination of expression patterns using fluorescent protein fusions expressed under the control of endogenous promoters showed that ALBA1, ALBA2 and ALBA4 are indeed expressed in mitotically active cells of root and leaf primordia, as is ECT2 (**Figure 3A,B**). The tight co-expression of ECT2 and ALBA proteins was also evident from analysis of published root single-cell mRNA-seq data^46,47^ (**Figure S4**). Further assessment of the subcellular localization by confocal microscopy indicated that ALBA1, ALBA2, ALBA4 and ALBA5 localize to the cytoplasm (**Figure 3C**), as do ECT2, ECT3 and ECT4^7,15,44^.

**Figure 3.**
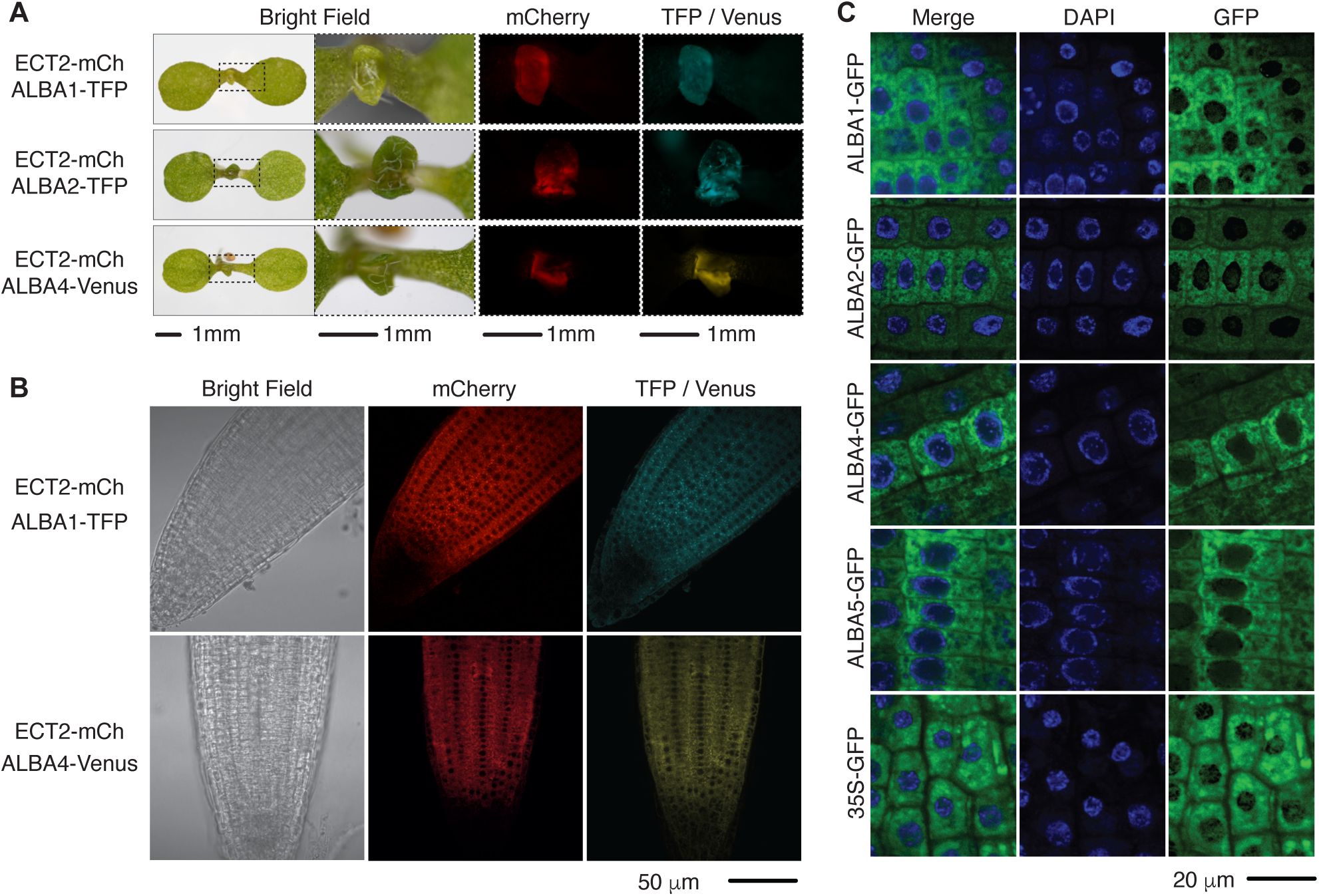
The expression patterns and subcellular localizations of ECTs and ALBAs overlap. **(A)** Fluorescence microscopy of 5-day old seedlings co-expressing ECT2-mCherry and ALBA1-TFP (top panel), ECT2-mCherry and ALBA2-TFP (middle panel), or ECT2-mCherry and ALBA4-Venus (bottom panel). **(B)** Confocal microscopy images of mCherry and GFP fluorescence in root tips of plants co-expressing ECT2-mCherry and ALBA1-TFP (top) or ECT2-mCherry and ALBA4-Venus (bottom). **(C)** Confocal images of GFP fluorescence and DAPI staining in root tips of plants expressing ALBA1-GFP, ALBA2-GFP, ALBA4-GFP, ALBA5-GFP and 35S-GFP.

The model further predicts that ALBAs and ECTs share a significant overlap in mRNA target sets, that they have juxtaposed binding sites around m^6^A sites in those target mRNAs, and that at least some direct mRNA targets associate less with ECTs *in vivo* in the absence of ALBA proteins. We previously demonstrated the feasibility of using TRIBE (Target Identification of RNA-binding Proteins by Editing)^48^ and iCLIP (Individual Nucleotide-Resolution Crosslinking and Immunoprecipitation)^49^ to address such predictions using transcriptome-wide analyses *in vivo*^28,44,50^. In TRIBE, the catalytic domain (cd) of the A-I RNA-editing enzyme ADAR is fused to the RNA-binding protein of interest, and targets are identified by mRNA-seq as mRNAs containing sites significantly more edited in cells expressing the ADAR_cd_ fusion compared to a free ADAR_cd_ control^28,48^. TRIBE can also be used to estimate differential protein-mRNA association between two conditions based on quantitative changes in editing proportions in target mRNAs. For example, many shared ECT2/3 target mRNAs are more highly edited by ECT3-ADAR_cd_ in the absence of ECT2, indicating that the two proteins compete for the same binding sites *in vivo*^44^. In iCLIP, target mRNAs are identified by co-purification with the protein of interest after covalent crosslinking *in vivo*, and binding sites are deduced from the position of frequent reverse transcription termination events at crosslink sites^49^. We therefore set out to test predictions on shared and interdependent ECT-ALBA target binding *in vivo* using combined iCLIP and TRIBE analyses focused on ALBA4 (long form), ALBA2 (short form) and ECT2.

### Identification of mRNA targets of ALBA4 using iCLIP

We first aimed to identify direct mRNA targets and binding sites of ALBA4 via iCLIP. To this end, we used transgenic lines expressing *ALBA4-GFP* under the control of the endogenous *ALBA4* promoter in the *alba4-1 alba5-1 alba 6-1* (henceforth, *alba456*) mutant background (**Figure S5A-B**), verified to carry T-DNA-induced knockout mutations in all three *ALBA* genes by RT-qPCR (**Figure S5C**) and western blot (**Figure S5D**) analyses. Initial immunoprecipitation tests with or without prior UV-crosslinking and followed by polynucleotide kinase (PNK) labeling established that RNA-protein complexes were specifically purified with ALBA4-GFP after UV crosslinking (**Figure 4A**). We therefore prepared and sequenced libraries from RNA immunopurified with ALBA4-GFP or GFP alone after crosslinking *in vivo* (**Figure S6A-D**), using the recently developed iCLIP2 protocol^51,52^. This effort identified 379,670 high-confidence replicated sites for ALBA4-GFP, corresponding to 7,744 genes (henceforth referred to as ALBA4 iCLIP2 targets). We further defined a "strong" set by filtering low scores, resulting in 63,695 sites mapping to 7,509 genes. In the GFP-only samples, only 81 sites in 13 genes were detected (**Figure 4B, Table S2**). Thus, nearly all ALBA4 iCLIP2 targets are strong candidates for *bona fide* ALBA4 target mRNAs.

**Figure 4.**
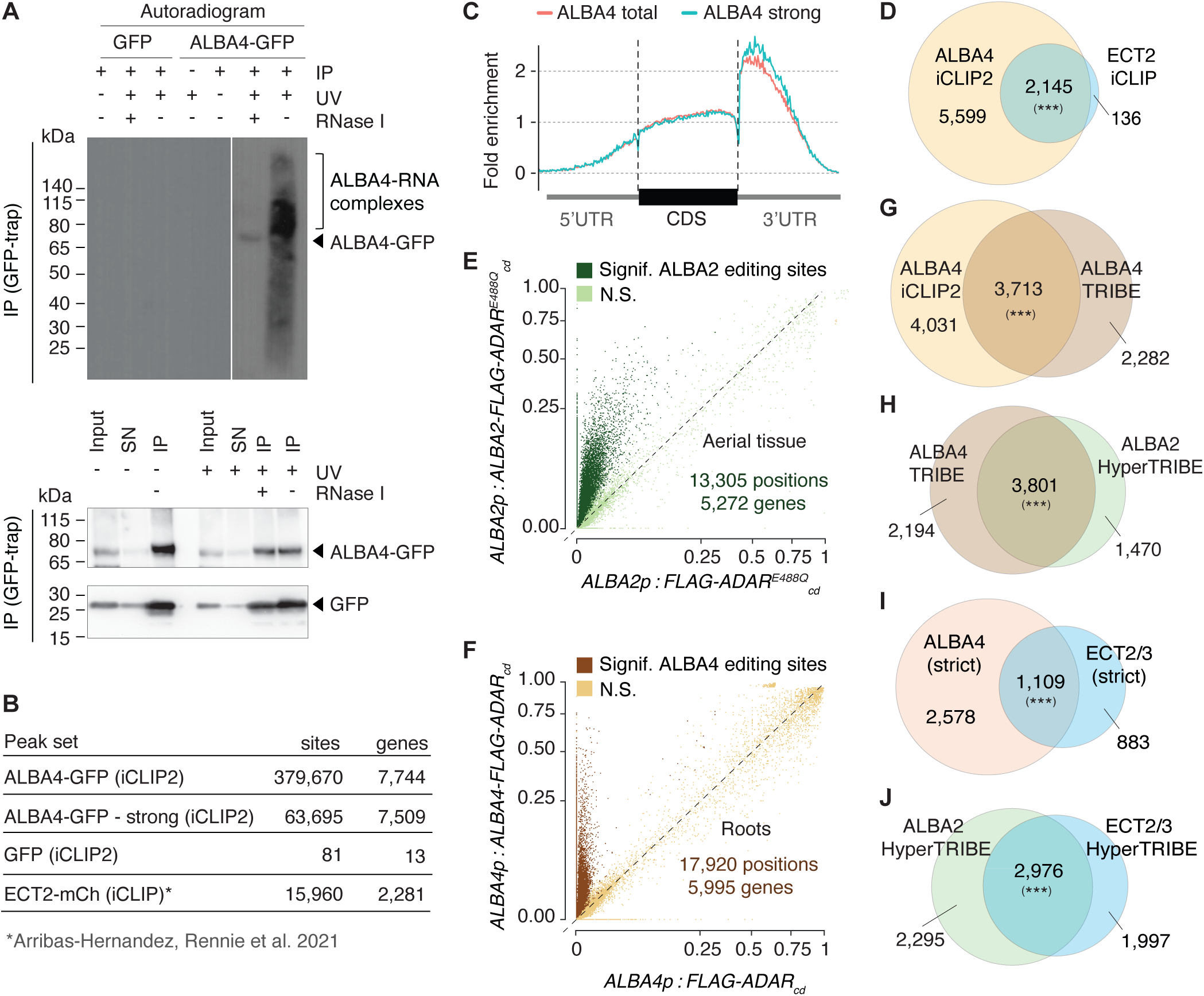
The mRNA targets bound by ECT2/3 and ALBA2/4 overlap substantially. **(A)** Top, autoradiogram of ^32^P-labelled RNA-protein complexes obtained by PNK/γ-^32^P-ATP labelling of immunopurified material from ALBA4-GFP- or GFP-expressing plants. Immunoprecipitations were carried out with or without UV crosslinking and after precipitation with GFP-Trap beads (IP+). (IP-) indicates mock immunoprecipitation with RFP-Trap beads. Treatment of the precipitate with RNase I (+ RNase) indicates the size of the precipitated protein. Marker positions and the location of the ALBA4-GFP RNA adducts are indicated. Bottom, immunoblots of input, supernatant (SN) after IP, and immunoprecipitated (IP) fractions, probed with GFP antibodies. Samples are pools of 3 independent lines for each genotype. **(B)** Number of called iCLIP peaks and associated genes for ALBA4-GFP, GFP alone and ECT2-mCherry^28^. Strong ALBA4-GFP peaks are defined as those with a score higher than the median, per gene. **(C)** Scaled metagene profiles showing the enrichment along the gene body (5’UTR, CDS or 3’UTR) of ALBA4-GFP iCLIP2 peaks. **(D)** Overlap of ECT2 and ALBA4 iCLIP mRNAs. The overlap is highly significant (p < 10^-16^, permutation test based on random sampling of genes from transcriptome with matched expression patterns, see Methods). **(E)** Scatterplot of the editing proportions (E.P.=G/(A+G)) of potential and significant editing sites (E.S.) determined by comparing mRNA-seq data obtained from transgenic lines expressing ALBA2-FLAG-ADAR or FLAG-ADAR in the Col-0 background, both under the control of the ALBA2 promoter (seedlings, shoot tissue). Significance was determined using the hyperTRIBER pipeline^57^, specifying an adjusted-p-value <0.01 and log_2_ fold-change > 1. **(F)** Same analysis as in (E), but carried out with roots of lines expressing ALBA4-FLAG-ADAR or FLAG-ADAR under the control of the ALBA4 promoter in the Col-0 background. **(G)** Overlap of ALBA4 targets identified using iCLIP2 and TRIBE analysis. The overlap is highly significant (p < 10^-16^, permutation test, as in D). **(H)** Overlap between ALBA4 TRIBE targets (roots) and ALBA2 HyperTRIBE targets (shoots). The overlap is highly significant (p < 10^-16^, permutation test, as in D). Most non-overlapping targets are expressed specifically in shoots or roots (**Figure S7**). **(I)** Overlap between high-confidence ALBA4 targets, supported by iCLIP and TRIBE, and ECT2/3 targets, supported by ECT2/3 HyperTRIBE and ECT2 iCLIP. The overlap is highly significant (p < 10^-16^, permutation test, as in D). **(J)** Overlap between ALBA2 HyperTRIBE targets and ECT2/3 HyperTRIBE targets. The overlap is highly significant (p < 10^-16^, as in D).

### mRNA target sets of ALBA proteins overlap significantly with those of ECT2/ECT3

We first noticed that ALBA4 iCLIP2 sites occurred in coding regions and, even more predominantly, in 3’-UTRs, with the 3’-UTR enrichment particularly apparent in the strong set (**Figure 4C**). Importantly, more than 90% of ECT2 iCLIP targets are also ALBA4 iCLIP2 targets (**Figure 4D**). Hence, ECT2 mainly binds to mRNAs that are also targeted by ALBA4. To corroborate this essential conclusion, we employed TRIBE to identify targets of both a long (ALBA4) and a short (ALBA2) ALBA protein family member by independent means. We used the improved variant HyperTRIBE relying on a hyperactive mutant of the ADAR_cd_^53^ for ALBA2, but had to proceed with TRIBE for ALBA4, because expression of the hyperactive ADAR_cd_ fused to ALBA4 was lethal (see Methods). In both cases, lines expressing comparable levels of free and ALBA-fused ADAR_cd_ were selected for mRNA-seq analysis (**Figure S7**). Significantly differentially edited sites between fused and free ADAR_cd_ exhibited higher editing proportions in the ALBA2/4-ADAR_cd_ fusions, as expected (**Figure 4E,F**). These differentially edited sites defined 5,272 target mRNAs for ALBA2 and 5,995 for ALBA4 (**Figure 4E,F, Table S3**). Using these target sets, ALBA4 iCLIP2 targets and the previously defined ECT2/3 targets^28,44^ for comparative analyses, we revealed the following three properties of ALBA2/4 and ECT2/3 target mRNAs and the relations between them. (1) The ALBA4 iCLIP2 target set is robust, because the overlap with ALBA4 TRIBE is significant (**Figure 4G**). In particular, TRIBE support of ALBA4 iCLIP2 targets is prominent for those target mRNAs with multiple called iCLIP peaks (**Figure S8A-B**). (2) ALBA4 and ALBA2 target a common set of mRNAs (**Figure 4H**) and differences between the two target sets can largely be explained by the tissue source used for the analysis (aerial tissues for ALBA2, roots for ALBA4) (**Figure S8C**). (3) The overlaps between the ECT2/3 target set and both the high-confidence set of ALBA4 targets supported by iCLIP2 and TRIBE and the set of ALBA2 HyperTRIBE targets are highly significant, as demonstrated by comparison to corresponding random target sets (**Figure 4I,J; Figure S8D-F, FigureS9, Table S4**). We conclude that ECT2/3 and ALBA2/4 mRNA target sets significantly overlap, thus fulfilling a second key requirement of the model of concerted mRNA binding by ECT-ALBA modules.

### ALBA proteins bind to pyrimidine-rich elements in the vicinity of m^6^A

We next analyzed positions of ALBA4 binding sites in their targets using the iCLIP2 data. Metagene analysis normalizing for region length showed a peak in the density of ALBA4 binding sites in 3’-UTRs, if less pronounced than ECT2 binding sites and m^6^A-sites, because ALBA4 binding sites also occur in coding regions as noted above (**Figure 5A**). The *RPS7A* and *TUBULIN ALPHA-5* genes provide illustrative examples of this close alignment of m^6^A, ECT2 and ALBA4 sites (**Figure 5B**). Both ALBA4 iCLIP2 and ECT2 iCLIP peaks^28^ are enriched upstream of m^6^A sites determined by Nanopore direct RNA sequencing^54^ (**Figure 5C,D**), with ALBA peaks situated either at or slightly upstream of ECT2 peaks (**Figure 5E**). Strikingly, the enrichment of ALBA4 peaks at m^6^A-sites was much more pronounced when considering peaks in ECT2 targets compared to non-targets. Indeed, the ALBA4 peak enrichment around m^6^A-sites in ECT2 non-targets showed a distribution similar to the location-matched background (**Figure 5F**). These key observations demonstrate that the important prediction of juxtaposition of ECT2 and ALBA4 binding sites on target mRNAs is fulfilled, and strongly suggest mutual dependence on target mRNA binding.

**Figure 5.**
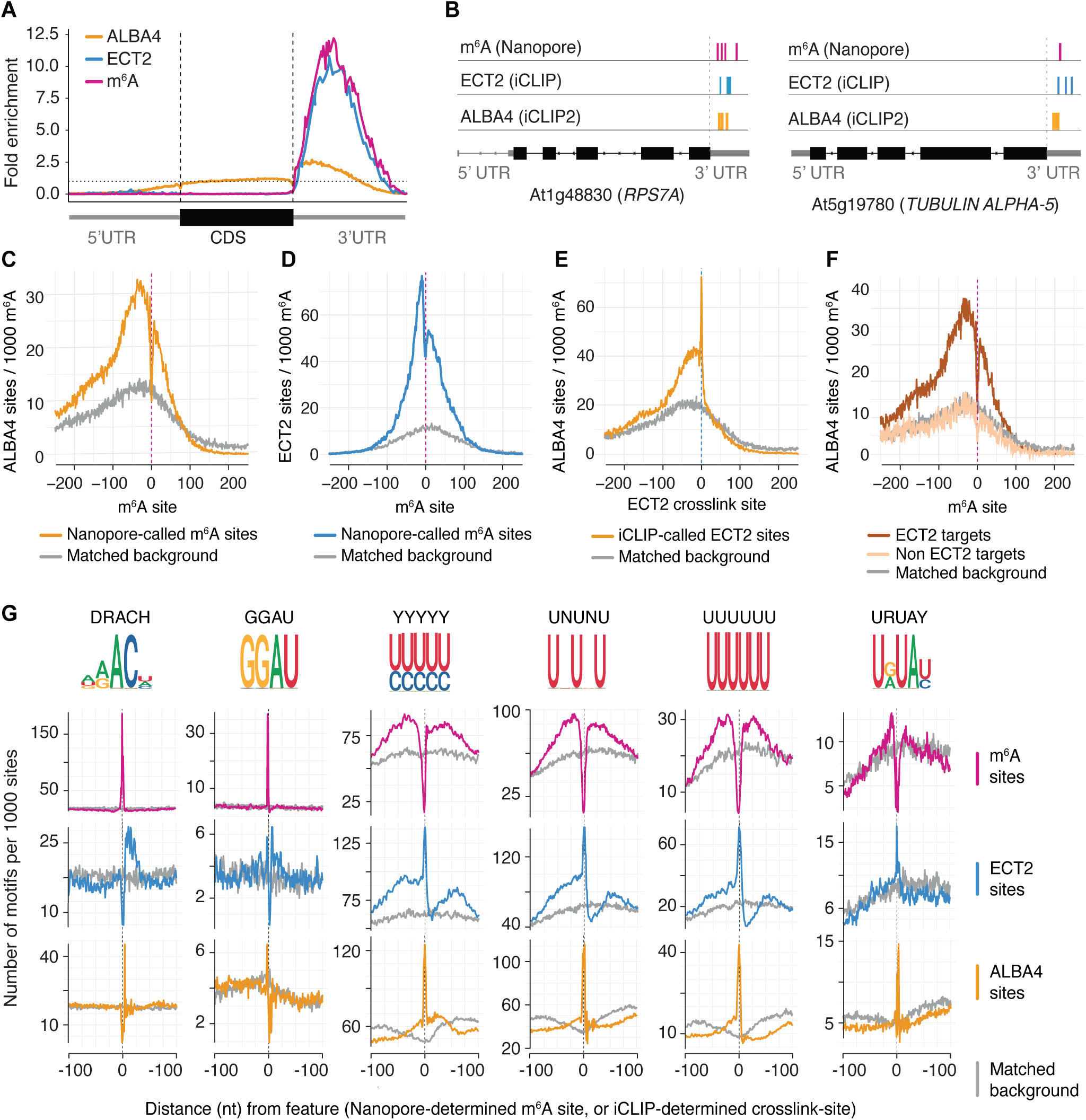
ALBA4 binds to pyrimidine-rich elements juxtaposed to m^6^A. **(A)** Scaled metagene profiles showing the enrichment along the gene body (5’UTR, CDS or 3’UTR) of the called ALBA4 iCLIP2 peaks. ECT2 iCLIP peaks^28^ and Nanopore-determined m^6^A density^54^ are shown for reference. **(B)** Representative examples of ECT2 and ALBA4 common targets showing the location of ALBA4 iCLIP2 and ECT2 iCLIP crosslink sites ^28^, and m^6^A sites^54^. **(C)** Number of ALBA4 iCLIP2 crosslink sites per 1000 Nanopore-derived m^6^A sites, as a function of distance from the m^6^A site. **(D)** Number of ECT2 iCLIP crosslink sites per 1000 Nanopore-derived m^6^A sites, as a function of distance from the m^6^A site. **(E)** Number of ALBA4 iCLIP2 crosslink sites per 1000 ECT2 crosslink sites, as a function of distance from the crosslink site. **(F)** Number of ALBA4 iCLIP2 crosslink sites per 1000 Nanopore-derived m^6^A sites, as a function of distance from the m^6^A site and according to whether containing genes are also targets of ECT2 or non-ECT2 targets. For each set, a matched background set was defined as positions on similarly expressed genes with a similar metagene distribution to the true set. **(G)** Number of the indicated motifs (selected from ^28^) per 1000 Nanopore-determined m^6^A sites (top), ECT2 iCLIP crosslink sites (middle) or ALBA4 iCLIP2 crosslink sites (bottom). For each set, a matched background set was defined as positions on similarly expressed genes with a similar meta-gene distribution to the true set.

Because we previously showed that several sequence motifs are enriched around ECT2 binding sites^28^, we went on to study whether any of these motifs were enriched at ALBA4 binding sites. We included 6 motifs identified as enriched around ECT2 iCLIP sites in our previous study^28^. This analysis revealed that uridine- or pyrimidine-rich motifs in the immediate vicinity of m^6^A/ECT2 binding sites are strongly enriched precisely at ALBA4 crosslink sites (**Figure 5G**), suggesting that these sequences may be ALBA4 binding sites *in vivo*.

### Deep learning supports pyrimidine-rich elements in the vicinity of m^6^A as determinants of ALBA4-ECT2 binding

One potential pitfall of this conclusion is that the photochemical properties of nucleobases result in a bias of UV-induced RNA-protein crosslinks to occur at uridines^55,56^ such that iCLIP sites can be located at nearby uridines if the actual binding site lacks this nucleotide. For example, many miCLIP sites obtained by UV crosslinking of an m^6^A-specific antibody to RNA *in vitro* map to uridines surrounding the uridine-depleted major m^6^A consensus site (DRACH)^28^. Therefore, we employed neural networks to identify sequence elements that distinguish m^6^A sites bound by ECT2/ALBA4 from m^6^A sites not bound by these proteins. We first collected Arabidopsis m^6^A sites from multiple published sources and curated a compendium of 41,883 high-quality, non-overlapping m^6^A sites which have properties highly consistent with the smaller set of sites identified by Nanopore direct RNA sequencing^54^ (**Figure S10**, **Table S5**, see Methods). Of these, 16,406 sites were annotated as ECT2-positive and 22,866 were ALBA4-positive (**Figure 6A**). We then used sequences surrounding all sites for input into a neural network trained simultaneously on two binary outputs: whether ECT2 was bound or unbound, and whether ALBA4 was bound or unbound (**Figure 6A**). This model performed well when predicting the presence of ALBA4 or ECT2 at m^6^A sites on gene sets excluded during model training (average AUC=0.74 (ECT2) and 0.76 (ALBA4), based on five-fold cross validation), with predicted binding probabilities clearly distinguishing between bound and unbound sites (**Figure 6B**). As expected, predicted binding probabilities for the two proteins correlated (PCC = 0.71, **Figure 6C**). Importantly, some differences between the two suggested that the model had learned specific sequence patterns relevant to each protein. To investigate this, we leveraged the filters learned in the first convolutional layer, since these represent motifs identified *de novo* by the model. We converted the sequences of the highest-scoring instances into position weight matrices (PWMs) and fit a generalised linear model predicting motif presence additively from the network-predicted ECT2 and ALBA4 binding probabilities (see Methods). From this model, the coefficient for each protein (motif score) can be interpreted as the effect of that protein controlling for the other (**Figure 6D**). This analysis identified the uridine-/pyrimidine-rich motifs UAUUUU and UUUACUUU as determinants of both ECT2-bound and ALBA4-bound m^6^A sites (**Figure 6D**). Indeed, the UAUUUU and UUUACUUU motifs were highly enriched at ALBA4 iCLIP sites and located just upstream of ECT2 iCLIP sites (**Figure 6E**), thus providing independent experimental evidence that these motifs act as ALBA4 binding sites. This conclusion is particularly important because it provides a simple molecular explanation for our previous machine learning-based finding that uridine- or pyrimidine-rich motifs are important for the distinction between m^6^A sites bound or not by ECT2^28^: juxtaposed m^6^A sites and uridine-/pyrimidine-rich elements provide the context required for binding of the ECT-ALBA module.

**Figure 6.**
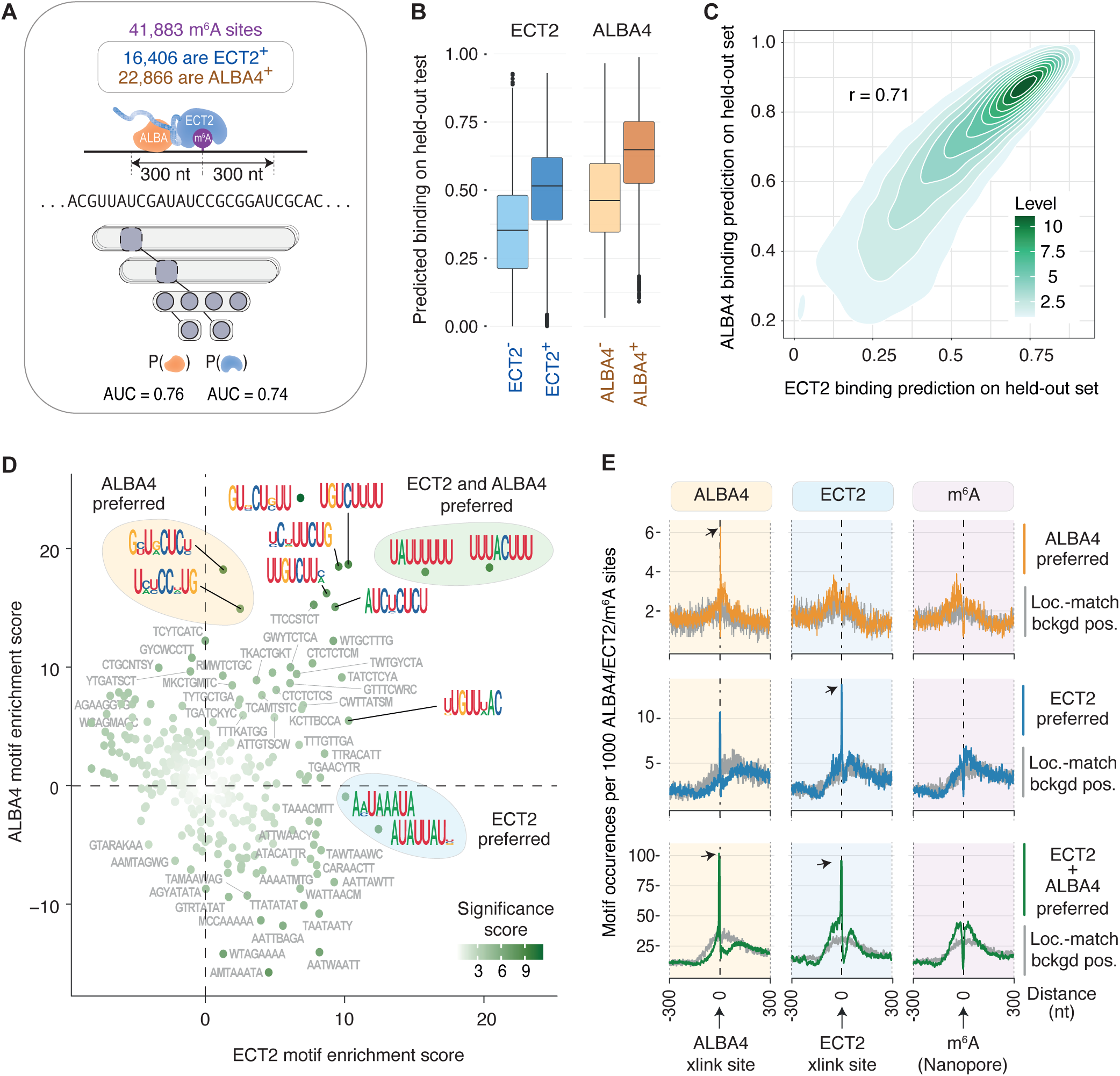
Neural network analysis identifies U-rich motifs in the vicinity of m^6^A as determinants of ALBA4-ECT2 binding. **(A)** Strategy for deep learning. m^6^A sites were annotated according to presence or absence of either ECT2 or ALBA4 and a convolutional neural network was trained which takes sequences surrounding m^6^A as input and predicts the probability of ECT2 and ALBA4 binding. **(B)** Boxplots showing predicted binding probabilities from the network, split according to protein and binding status. **(C)** Scatter plot of the predicted ALBA4 binding probabilities against the ECT2 binding probabilities from the network. Counts depict the density of sites. **(D)** Output-specific enrichment scores for *de novo* motifs learned by convolutional neural network, calculated using a generalized linear model for predicting motif presence from predicted presence of ECT2 and ALBA4 at m^6^A-centered sequences using model. Colored circles indicate interesting motifs determined as specific to ALBA4 (yellow), ECT2 (blue) or both (green). **(E)** Enrichment of motif sets indicated in D around ALBA4 iCLIP2, ECT2 iCLIP and Nanopore-derived m^6^A sites ^54^. Grey shows location-matched background positions.

### Binding to target mRNA in vivo involves mutual ALBA-ECT dependence

We next assessed whether ALBA proteins are necessary for mRNA target association of the wild type ECT2 protein. Initially, we used the CLIP-PNK assay with ECT2-mCherry expressed in wild type, or the *alba1-2 alba2-2 alba4-1 alba5-1* (henceforth *alba1245*) or *alba456* mutant backgrounds, carrying T-DNA insertion alleles in the corresponding *ALBA* genes (**Figures S2B and S5**, see Methods). These experiments showed that ECT2-mCherry associated with less RNA in the *alba* mutants compared to wild type, with the clearest effects (∼2.5-fold reduction) observed in *alba456* mutants (**Figure 7A, Figure S11A**). We next used ECT2 HyperTRIBE to estimate the relative target mRNA binding in wild type and in *alba1245* mutants by differential editing. We chose this method both to gain sensitivity and to assess directly whether mRNAs that associate less with ECT2 *in vivo* in *alba1245* mutants are in fact dual ECT2/ALBA targets. We selected five independent lines expressing ECT2-ADAR in both wild type and *alba1245*, and performed mRNA-seq of root tissues to provide the raw data for analysis of differential editing. Positions exhibiting significant differential editing according to the hyperTRIBER package^57^ were strongly biased in the direction of lower editing in *alba1245,* although these results were potentially biased by the expression of the ECT2-ADAR fusion protein not being balanced between the two conditions (**Figure S11B,C,H**). For this reason, we developed a highly robust alternative statistical modelling approach, correcting the editing proportions for mRNA levels of ADAR and to obtain a smaller, high confidence set of significantly differentially edited sites between the two backgrounds (Methods). As a further control, we also performed differential editing analysis using only those replicates whose ECT2-ADAR expression was nearly perfectly matched as judged by both mRNA-seq read densities and protein blots, resulting in a smaller set of sites which overlapped significantly with the set from the robust modelling approach (**Figure S11F,G**). Overall, these analyses converged on the same conclusion: editing proportions in ECT2/ALBA4 mRNA targets tended to be higher in wild type than in *alba1245* mutants, indicating that ALBA proteins facilitate target mRNA binding of ECT2 *in vivo* (**Figure 7B,D**). Because the structural model of the ALBA-ECT2 interaction suggests that RNA association by the ALBA domain may also be enhanced by ECT proteins, we did the reciprocal experiment with the short ALBA2 protein. Thus, we expressed ALBA2-ADAR either in wild type or *ect2-3 ect3-2 ect4-2* (*Gte234*) mutant backgrounds and carried out analysis of differential editing proportions as above. We found that editing proportions of ALBA2-ECT2/3 targets were higher in wild type than in *Gte234* mutants (**Figure 7C,E, Figure S11D,E,I**), indicating that there is mutual ALBA-ECT dependence for mRNA target association *in vivo*. Taken together, our TRIBE-based assessment of target mRNA association *in vivo* supports the conclusion that the ALBA domain acts as a unit with the YTH domain to facilitate m^6^A-binding.

**Figure 7.**
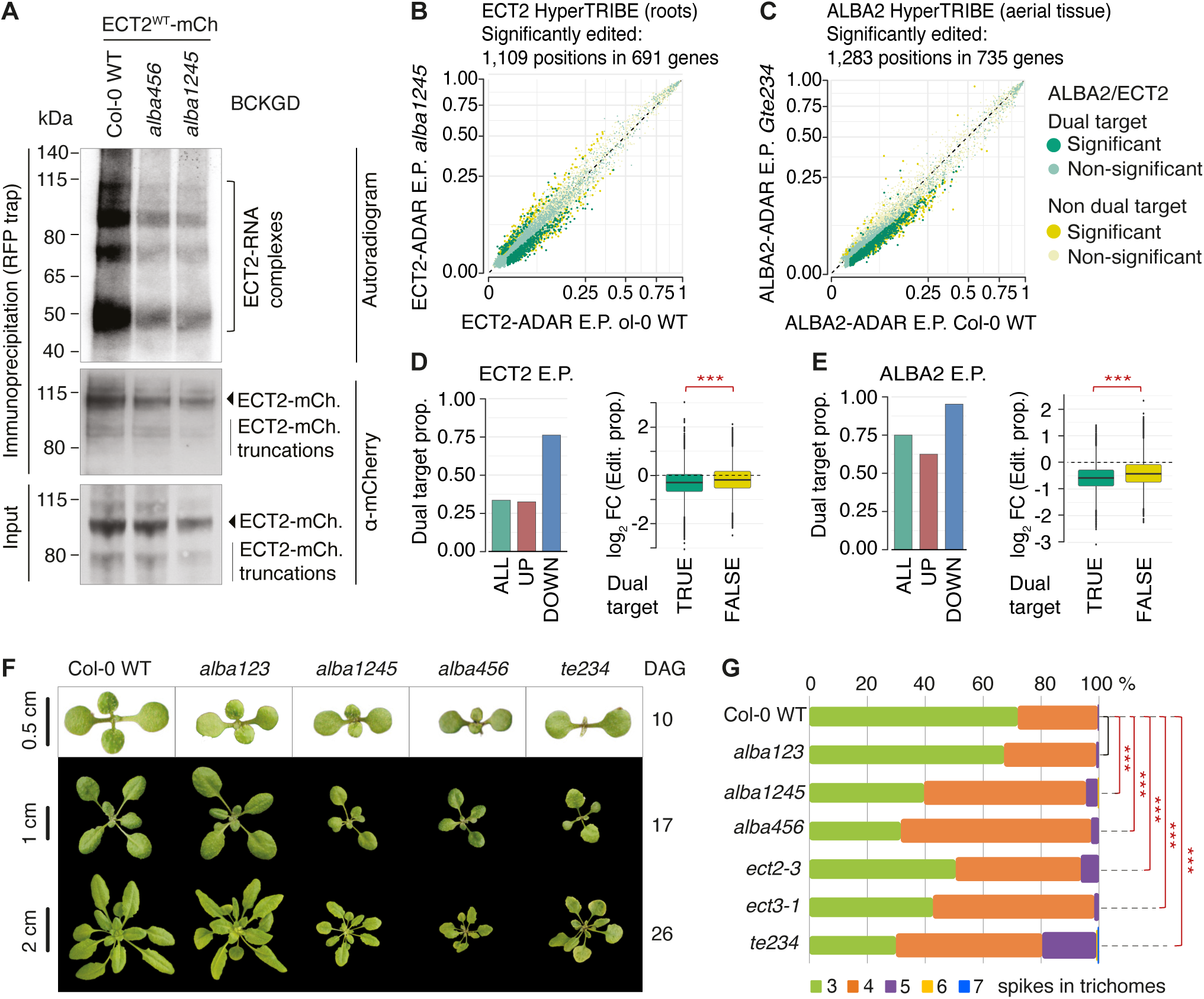
ALBA proteins are required for ECT2 target mRNA binding and biological function. **(A)** Results of an *in vivo* UV crosslinking-ECT2-mCherry immunoprecipitation experiment, followed by PNK-labelling of precipitated RNA with γ-^32^P-ATP. Top panel, autoradiogram of ^32^P-radiolabelled RNA-protein complexes purified from plants expressing ECT2^WT^-mCherry in the indicated genetic backgrounds. Molecular weight marker positions and the location of the verified ECT2-mCherry-RNA complexes^28^ are indicated. The presence of several bands of unequal intensity is due to partial proteolysis of the ECT2 IDR during immunoprecipitation, and differential labelling efficiency of the different RNPs^28^. Middle and bottom panels, mCherry immunoblots of the immunoprecipitated (middle) and total fractions (input, bottom). Samples were pools of 3 independent lines for each genotype. **(B)** Scatter plot showing the editing proportions of ECT2-ADAR-catalyzed editing sites between Col-0 WT and *alba1245*. Green, sites whose change in editing proportions is statistically significant and that are located in dual-bound mRNAs. Yellow, sites whose change in editing proportions is statistically significant but that are located in mRNAs not targeted by both ECT2 and ALBA4 (non-dual bound). Light green/light yellow, candidate sites whose change in editing proportions is not statistically significant. **(C)** Scatter plot showing the editing proportions of ALBA2-ADAR-catalyzed editing sites between Col-0 WT and *ect2-3 ect3-2 ect4-2* (*Gte234*). Color scheme as in (B). **(D)** Quantification of the tendency of sites differentially edited by ECT2-ADAR between Col-0 and *alba1245* to be less highly edited in *alba1245*. Left, histogram showing the fraction that sites in dual-bound targets comprise of either less highly edited sites in *alba1245* (down) or more highly edited sites in *alba1245* (up). The histogram also illustrates the fraction of editing sites in dual-bound targets relative to all editing sites for comparison. Right, boxplot showing the median log_2_ differential editing proportions for editing sites either in dual-bound mRNA targets (true) or in other mRNAs (false). Asterisks indicate p-values from 2-sample t-test: ***p < 0.001. **(E)** Quantification of the tendency of sites differentially edited by ALBA2-ADAR between Col-0 and *Gte234* to be less highly edited in *Gte234*. Analogous to the analyses presented in (D) for ECT2-ADAR in Col-0 vs *alba1245*. Asterisks indicate p-values from 2-sample t-test: ***p < 0.001. **(F)** Representative photographs of seedlings and rosettes of the indicated genotypes at three different time points given in days after germination (DAG) in soil. **(G)** Trichome branching sorted by number of spikes in the indicated genotypes. Branches were counted on at least 150 trichomes on each of at least 6 plants for each genotype (*n* = ∼1000). Data were fitted to a proportional odds model in R for statistical analyses (see Methods). Asterisks indicate Bonferroni-corrected p-values: ***p < 0.001. Black bar indicates no significant difference.

### Inactivation of ALBA and ECT genes cause similar developmental phenotypes

We finally characterized mutants in *ALBA* genes to assess whether they exhibit phenotypes characteristic of reduced m^6^A-ECT function. As previously reported, single *alba* mutants (**Figure S2**) did not show obvious developmental phenotypes^43^. In contrast, *alba123* mutants with lesions in all three ALBA-only-encoding genes, and in particular *alba1245* and *alba456* mutants showed pleiotropic developmental defects including slower growth, defects in leaf morphology and delayed flowering (**Figure 7F, Figure S12A,B**). Similar observations on smaller stature of *alba456* mutants have been reported by others^43^. Although some of these phenotypes are reminiscent of phenotypes displayed by *ect2 ect3 ect4* mutants, they are not identical. We therefore assessed a quantifiable phenotype seen consistently in mutants with defects in m^6^A-ECT function: branching of leaf epidermal hairs (trichomes, ^58^) where defects can be detected even in single *ect2* and *ect3* mutants^7,13,59^. We found that *alba1245* and *alba456* mutants showed increased trichome branching (**Figure 7G**), with a phenotypic strength intermediate between single mutants in *ect2* or *ect3* and the *te234* triple knockout mutant. Importantly, ECT2 protein levels in *alba1245* and *alba456* mutants were only slightly lower than in wild type (**Figure S12C**), excluding the trivial possibility that the phenotypic similarity between composite *alba* and *ect* mutants is due to drastically reduced ECT protein levels in *alba1245* and *alba456*. We conclude that the developmental defects of composite *alba* mutants are consistent with defective m^6^A-ECT function, as predicted by the model of m^6^A-ECT interaction facilitated by ALBA proteins.

## DISCUSSION

Our results on ALBA-ECT interaction and target binding *in vivo* provide strong support for the conclusion that the YTH domain of major plant YTHDF proteins is insufficient for full m^6^A binding *in vivo*, because it requires facilitation by ALBA proteins. In the following paragraphs, we discuss how this new understanding of the m^6^A-YTH interaction impacts the thinking of m^6^A-mediated genetic control in plants and other eukaryotes.

### Functional implications of recognition of m^6^A by the ALBA-YTHDF module

The discovery that m^6^A reading in plants involves YTHDF-m^6^A binding modulated by a third player, the ALBA proteins, introduces increased potential to integrate information into combinatorial control of biological effects of m^6^A. A key determinant of those effects is the fraction of m^6^A target mRNAs bound by YTHDF, in turn determined by the stoichiometry of m^6^A modification in mRNA, and YTHDF concentration and affinity for m^6^A-sites. Since we now understand that the affinity is not a constant, but must be tunable via, for instance, ALBA concentration and modification, we envision that plants have evolved to take advantage of this combinatorial potential to generate a gradient of m^6^A outputs that matches the cellular environment measured by multiple environmental and developmental sensors.

### Conservation of the ALBA-YTHDF unit and generality of RBP-assisted m^6^A-YTH interaction

It is an important observation that Arabidopsis YTHDF proteins both with and without the molecular properties required to complement organogenesis defects of *te234* mutants ^20^ retain the conserved YAIM and interact with ALBA4. This observation further supports the generality of ALBA-assisted m^6^A-binding among Arabidopsis YTHDF proteins. Thus, it is a pertinent question how widespread this phenomenon is. The YAIM is deeply conserved in land plant YTHDF proteins, strongly suggesting that the ALBA-YTHDF unit is conserved over the 500 My of land plant evolution. Beyond land plants, the YAIM is not conserved, and fungal and animal ALBA-family proteins are so divergent in sequence that conservation of the details of their molecular functions cannot be assumed. In addition, *Trypanosoma brucei* where ALBA proteins clearly perform functions in mRNA control^39,40^ does not encode YTHDF proteins, providing an example that the two families do not always have linked functions in eukaryotes. These observations raise two immediate questions.

First, given the deep conservation of the YTH domain, it is of interest how the m^6^A-YTHDF interaction is made efficient in organisms where ALBA proteins are unlikely to assist binding directly as in plants. We see two possible answers. Either other, as yet unidentified classes of RBPs evolved to facilitate m^6^A reading by YTHDF proteins, or the YTHDF proteins evolved to read m^6^A independently of other RBPs. In the latter case, comparative structure-function studies between, for instance, Arabidopsis and human YTHDF-m^6^A-RNA interactions should reveal the probably subtle structural features that may allow ALBA-independent efficient m^6^A-interaction. In this context, an YTH-proximal element in the IDRs of mammalian YTHDF proteins is of particular interest for at least two reasons. First, its location relative the YTH domain is reminiscent of the YAIM described here for plant YTHDFs. Second, it is predicted by AlphaFold^60^ to engage in YTH-domain interactions, perhaps via disorder-to-order transition upon RNA-binding to stabilize the RNA-bound form^61^, as observed for the *Schizosaccharomyces pombe* YTH-domain protein Mmi1^62^. The existence of non-ALBA facilitators of YTHDF-m^6^A binding in other organisms should not be entirely discarded, however. The mammalian IGF2BP/IMP/ZBP family of RBPs has been suggested to act as m^6^A readers based on multiple lines of evidence, including m^6^A-dependent target mRNA association and the similar positions of m^6^A sites and IGF2BP2 CLIP sites in 3’-UTRs of target mRNAs^63^. Because the m^6^A mapping methodology used at the time had limited resolution, it is possible that m^6^A sites are in fact adjacent to IGF2BP2 CLIP sites, particularly since the IGF2BP/IMP/ZBP recognition element (CAUH) defined in previous transcriptome-wide studies^64^ is not identical to the DRACH m^6^A consensus site. The slight off-set between IGF2BP CLIP site and m^6^A distributions^63^ is indeed reminiscent of the 3’-UTR distributions of m^6^A sites and ALBA4 iCLIP sites observed here, and the identification of IGF2BP2 as a prominent interactor of YTHDF1/2/3 in IP-MS experiments^65^ is more easily reconciled with a function in facilitated m^6^A binding by YTHDFs than direct m^6^A binding competing with YTHDFs. Thus, in light of our results on the ALBA-YTHDF-m^6^A module in plants, it may be appropriate to consider whether facilitated m^6^A-reading by YTHDF proteins could have evolved independently in several eukaryotic lineages, and, for mammals in particular, whether a function as a facilitator of m^6^A reading might explain many of the results originally interpreted to reveal a direct reader function of the IGF2BPs^63^.

Second, which molecular functions do ALBA proteins fulfill independently of YTHDF proteins? Such functions are anticipated for a number of reasons. First, while most ALBA4 mRNA binding sites in 3’-UTRs appear to be linked to m^6^A sites, binding sites in open reading frames were even more numerous and were found in mRNAs with no evidence of m^6^A modification or ECT2/3 binding. Indeed, ALBA proteins have been found to play a role in heat adaptation via regulation of Heat Shock Factor-encoding mRNAs, primarily with binding sites in open reading frames^43^. Second, even the YTHDF-linked ALBA functions may involve properties in addition to assisted m^6^A-binding, because many ECT2-associated proteins were depleted in the immunoaffinity-purified fraction of the ECT2^YAIM^ mutant defective in ALBA interaction. Finally, we note that while this report identifies a molecular role of the ALBA domain, it does not address the function of the C-terminal IDR of long ALBA proteins, expected to be of considerable biological importance given the stronger phenotypes of *alba456* compared to *alba123* mutants, as reported here and by others^43^.

## METHODS

### Plant material and growth conditions

All lines used in this study are in the *Arabidopsis thaliana* Col-0 ecotype. The following mutant and transgenic lines mentioned have been previously described: *ect2-1 ect3-1 ect4-2* (*te234*)^7^, *ect2-1 ECT2^W464A^-mCherry*^7^, *ect3-2 ECT3-Venus*^7^, *ect2-1 HA-ECT2* ^21^. The *alba1-1 (*GABI_560B06), *alba1-2* (SALK_069210), *alba2-1* (GABI_128D08), *alba2-2* (SALKseq_069306), *alba3-1* (SAIL_649_E11), *alba4-1* (SALK_015940), *alba5-1* (SALK_088909) and *alba6-1* (SALK_048337) single mutants were obtained from the Arabidopsis Biological Resource Center (ARBC). Seeds were sterilized by immersing them in 70% EtOH for 2 min, followed by incubation in 1.5% NaOCl, 0.05% Tween-20 for 10 min, after which the seeds were washed twice with H_2_O. The seeds were then spread on plates containing Murashige & Skoog (MS) medium (4.1 g/l MS salt, 10 g/l sucrose, 8 g/l Bacto agar). The plates were stratified in darkness at 4°C for 2–5 days before transfer to Aralab incubators at 21°C, with a light intensity of 120 μmol/m^2^ and a photoperiod of 16 h light/8 h dark. When needed, after 10 days of growth, seedlings were transferred to soil and kept in Percival incubators under identical settings.

#### Generation of *ect2-5 ect3-1 ect4-2* by CRISPR-Cas9 genome engineering

For the targeted creation of an in-frame deletion mutant at the endogenous ECT2 locus, we employed the pKIR1.1 CRISPR-Cas9 system^70^. Two plasmids, pKIR1.1-ect2-N8A and pKIR1.1-ect2-N8B, expressing sgRNAs were constructed by ligating oligonucleotides that target *ECT2* into pKIR1.1, as described^70^. The crRNAs were designed to yield a deletion resembling ECT2^ΔN8^ as closely as possible. The plasmids were then transformed into *ect3*-*1 ect4*-*2* mutants, and transformants were selected on MS-agar supplemented with 25 μg/mL hygromycin. After transfer to soil, plants with deletions in *ECT2* were identified via PCR using primers spanning the deletion. Progeny from plants with deletions of the expected size, as confirmed by migration in a 1% agarose gel, were plated on MS supplemented with 25 μg/mL hygromycin. Hygromycin-sensitive plants, indicative of the absence of Cas9 and homozygosity of the deletion, were rescued and transferred to MS-agar for recovery. Subsequently, these plants were genotyped and Sanger sequenced for identification of in-frame deletions. Western blotting, utilizing antibodies raised against synthetic peptides in the ECT2 IDR outside the deleted region^7^, was performed to confirm the in-frame deletion. Primers are listed in Table S6.

### Construction of transgenic lines

To generate the constructs *pro(ALBA2):ALBA2-FLAG-TFP:ter(ALBA2)*, *pro(ALBA4):ALBA4-VENUS:ter(ALBA4)*, *pro(ALBA2):ALBA2-FLAG-ADAR:ter(ALBA2)*, *pro(ALBA4):ALBA4-FLAG-ADAR:ter(ALBA4)*, *pro(ECT2):ECT2^YAIM^-mCherry:ter(ECT2),* PCR-amplified DNA fragments were pieced together by USER cloning^71^ in all cases except for *pro(ECT2):ECT2^YAIM^-mCherry:ter(ECT2)* in which an appropriate dsDNA containing the YAIM-mutations was synthesized (Integrated DNA Technologies, gBlocks). As template for PCR, we used plasmids containing wild-type *pro(ECT2):ECT2-mCherry:ter(ECT2)*^7^ for *ECT2-mCherry* constructs, *pro(ECT2):ECT2-FLAG-ADAR:ter(ECT2)* for *FLAG-ADAR* constructs^28^, and *pro(ECT3):ECT3-VENUS:ter(ECT3)* for *VENUS* constructs^7^. DNA fragments were amplified using dU-substituted primers and KAPA Hifi Hotstart Uracil+ ReadyMix^71^. The amplified fragments were inserted into the pCAMBIA3300-U vector, a modified version with a double PacI USER cassette^72^. To clone *pro(ALBA1):ALBA1-FLAG-TFP:ter(ALBA1)*, we made use of Greengate cloning. Briefly, PCR fragments were amplified using Thermo Scientific Phusion High-Fidelity DNA Polymerase (NEB) and ligated into entry vectors through BsaI-restriction cloning. The *pro(ALBA1):ALBA1* gDNA fragment was subcloned into pGEM-T Easy by A-tailing (Promega) prior to BsaI-restriction cloning. The vectors containing *pro(ALBA1):ALBA1* (in pGEM-T Easy), linker-*TFP* (pGGD003), *ALBA1* 3’UTR and downstream sequences (in pGGE000), and the D-AlaR cassette (pGGF003) were combined in a ’Greengate reaction’ using BsaI-HF (NEB), T4 DNA-Ligase (Thermo Scientific), and pGGZ001 as the destination vector. *pro(ALBA2):ALBA2-FLAG-TFP:ter(ALBA2),* and *pro(ALBA4):ALBA4-FLAG-Venus:ter(ALBA4)* fusions were constructed by USER cloning with the primers listed in **Table S6**. To clone *ALBA1-GFP, ALBA2-GFP, ALBA4-GFP,* and *ALBA5-GFP* used for confocal microscopy, we employed Gateway cloning. *ALBA* gene fragments, including 5’-regions, exons/introns to the gene’s end (excluding the stop codon), were amplified with attB1 and attB2 sites for Gateway cloning using KOD Hot Start DNA Polymerase. Purified amplicons were cloned into pDONR/Zeo via Gateway BP Clonase II (Thermo Fisher) and transformed into *E. coli* α-select cells. Subsequently, entry clones were recombined with the destination vectors pMDC111 and pMDC164, respectively (^73,74^ via Gateway LR Clonase II (Thermo Fisher) to generate expression clones. All plasmids were verified through restriction digestion and sequencing before being transformed into respective plants using Agrobacterium-mediated floral dip^75^. Primers are listed in **Table S6**.

### Screening for *te234* complementation

Screening of primary transformants (T1s) expressing wild-type, deletion or point mutant variants of ECT2-mCherry in the *te234* background was done as previously described ^21^. In brief, primary transformants were selected on MS-agar plates containing glufosinate ammonium (7.5 mg/L (Sigma)) to select plants with the transgene and ampicillin (10 mg/l) to restrict agrobacterial growth. Nine days after germination, primary transformants were categorized according to the size(s) of the first true leaves: full complementation (*s*L≥L1 mm), partial complementation (0.5 mmL<L*s*L<L1 mm), or no complementation (*s*L≤L0.5 mm). The complementation percentages were then determined by dividing the number of seedlings in each complementation category by the total number of transformants.

### Statistical analysis of complementation data

Statistical significance of the different T1 complementation categories was determined using Fisher’s exact test, and the Holm–Bonferroni method was applied to address multiple testing. Student’s t-test was used to evaluate the significance of differences in leaf size between Col-0 WT, *de34* (*ect3-1 ect4-2*), *te234* (*ect2-3 ect3-1 ect4-2*), and the CRISPR-generated *ect2-5 ect3-1 ect4-2*.

### Analysis of trichome phenotypes

Counts of trichomes with different numbers of branches and the statistical analysis of the raw data were done as described^7^.

### Western blotting

Western blotting was performed as described ^21^. In brief, 100-300 mg of tissue were ground in liquid nitrogen and resuspended in 5 volumes of IP buffer (50 mM Tris–HCl pH 7.5, 150 mM NaCl, 10% glycerol, 5 mM MgCl2, and 0.1% Nonidet P40), supplemented with 1x protease inhibitor (Roche Complete tablets) and 1 mM DTT. The lysate was centrifuged at 13,000 g for 10 min and 4× LDS sample buffer (277.8 mM Tris–HCl pH 6.8, 44.4% (v/v) glycerol, 4.4% LDS, and 0.02% bromophenol blue) was added to a final concentration of 1× LDS. Subsequently, the samples were denatured at 75°C for 10 min and run on a 4–20% Criterion™ TGX™ Precast gel in 1× Tris-glycine, 0.1% SDS buffer at 90–120 V for ∼1 h on ice. The proteins were transferred onto an Amersham Protran

Premium nitrocellulose membrane (GE Healthcare Life Sciences) in cold transfer buffer (1 ×LTris-glycine, 20% EtOH) at 80 V for 1 h on ice. The membrane was then blocked in 5% skim milk in PBS-T (137 mM NaCl, 2.7 mM KCI, 10 mM Na_2_HPO_4_, 1.8 mM KH_2_PO_4_, pH 7.4, 0.05% Tween-20) for 30 min. After blocking, membranes were probed with antibodies specific for ECT2^7^, ALBA1 (1:2000, see below), ALBA4 (1:1000, see below), or commercially available antibodies against mCherry (Abcam ab183628, 1:2,000 dilution) at 4°C over night. Membranes were then washed three times in PBS-T, incubated with HRP-coupled goat-anti-rabbit antibody and developed using chemiluminescence detection, as previously described^7^.

### RNA extraction and qRT-PCR

Total RNA was extracted from frozen and ground plant powder using TRIzol® (1 mL per 500 mg sample). 14 μg RNA was treated with 14 μL of RQ1 RNase-Free DNase (Promega) and 1 μL of RNaseOut^TM^ Recombinant RNase Inhibitor (Invitrogen) following the manufacturer’s protocol. The RNA was then purified using the QIAgen RNeasy mini kit following the RNeasy column clean-up protocol. The RNA quantity and quality were determined via NanoDrop and agarose gel electrophoresis. cDNA was prepared using SuperScript® III Reverse Transcriptase (Invitrogen), with the addition of RNaseOut^TM^. For qRT-PCR, 0.4 μL 10 μM specific primer pairs (mixture of forward and reverse primers) was mixed with 10 μL SensiFAST SYBR (Bioline) mastermix and 9.6 μL of cDNA. All the qRT-PCR reactions were performed in three technical replicates, carried out on a QIAGEN Rotor-Gene-Q real-time PCR machine and analyzed with the Rotor-Gene 6000 series software (QIAGEN). *CYCLOPHILIN* (At2g29960) was used for normalization. Primers are listed in **Table S6**.

### CLIP-PNK assays of ECT2-mCherry variants

12-day-old seedlings were UV-crosslinked with 2000 mJ/cm^2^ and ground into a fine powder in liquid nitrogen. Immunoprecipitation with RFP-trap beads (Chromotek), washes, DNase and RNase digestion, PNK labelling, SDS-PAGE, membrane transfer and autoradiography were performed as described in^28^. We used 20 μL of beads for 1 g of tissue in 1.5 mL of iCLIP buffer for every sample.

### Immunoprecipitation and LC-MS

Immunoprecipitations of ECT1-TFP, ECT2-mCherry variants and ECT3-Venus were performed as described by^21^, while immunoprecipitations of ALBA4-GFP or GFP were performed as described by^76^. Briefly, 7-day-old seedlings expressing ALBA4-GFP or GFP alone were harvested and ground into fine powder using liquid nitrogen. For each replicated, 0.5 g of ground plant tissue was homogenized in 1.5 mL IP buffer (50 mM Tris-HCl pH 7.5, 150 mM NaCl, 10% glycerol, 0.1% Triton-X100) supplemented with 2% (w/v) PVP40, Roche Complete Protease Inhibitor cocktail (1 tablet/50 mL), 100 μM MG132, 1 mM PMSF and Sigma Plant Protease Inhibitor cocktail (1/30 v/v). Samples were centrifuged at 16,000 x g for 5 min at 4°C, the supernatant was transferred to a new tube and centrifugation was repeated for 10 min. The supernatant was again transferred to a new tube and filtered through a 0.45 μm filter. For Co-IP, 1 mL of cell extract at a concentration of 2 μg/μL was first added to 50 μL of sepharose beads for pre-clearing and incubated for 30 min at 4°C with constant rotation. After centrifugation at 1000 x g for 1.5 min at 4°C, the cell extract was added to 20 μL GFP-Trap beads and incubated for 2.5 h at 4°C with constant rotation. The beads were washed 4x in Co-IP wash buffer (50 mM Tris-HCl pH 7.5, 150 mM NaCl, 10% glycerol, 0.05% Triton-X100, Roche Complete Protease Inhibitor cocktail (1 tablet/50 mL)) and proteins were eluted by addition of 40 μL 2x LDS sample buffer to the beads and incubation at 70°C for 10 min. For control samples treated with nucleases, beads were washed once in Co-IP wash buffer (+ 10 mM MgCl_2_) after the IP. Beads were then resuspended in 100 μL Co-IP wash buffer (+ 10 mM MgCl_2_) and treated with 2 μL Turbo DNase (Thermo Fisher Scientific) and, optionally, 5 μL of a 1:50 dilution of RNase I (Ambion) for 10 min at 37°C and 1200 rpm. Beads were then washed three times with Co-IP wash buffer and elution was performed as described above. Mass spectrometry data was analysed as in^21^ .

### Protein expression of ALBA1

An ALBA1 (AT1G29250) cDNA was amplified from oligo(dT)-primed reverse transcription products of DNase-treated total RNA from Col-0 wild type using the primer set MT303- MT304.The resulting PCR product was ligated in frame downstream of His_6_-SUMO in pET-24-derived vector containing His_6_-SUMO (Twist Bioscience). For recombinant protein expression, the plasmid encoding His-SUMO-ALBA1 was transformed into *E.*LJ*coli* BL21 (DE3 7tRNA) codon plus. Cells were grown at 37°C in LB medium supplemented with 35 μg/ml kanamycin, and expression was induced at OD_600_L≈L0.6 by addition of 0.5 mM IPTG. Following induction, the cells were grown at 18°C overnight and harvested by centrifugation. The cell pellet was resuspended in 20 mM Tris–HCl (pH 8), 10 mM imidazole, and 300 mM NaCl supplemented with 1 mM DTT and EDTA-free protease inhibitor (cOmplete; Roche). Cells were lysed once using a French press (20,000 psi). Crude lysate was cleared by centrifugation at 30,000 *g* for 30 min at 4°C and filtered through a 0.45-μm membrane. His-SUMO-ALBA1 was purified on Ni^2+^-NTA resin by incubation for 1 h at 4°C after which the beads were washed in wash buffer (20 mM Tris–HCl pH 8, 20 mM imidazole, 200 mM NaCl), and the bound protein was eluted in elution buffer (300 mM imidazole, 20 mM Tris–HCl pH 8, 300 mM NaCl). The eluted protein was dialysed overnight into 20 mM Tris-HCl pH 8, 200 mM NaCl, 1 mM 2- mercaptoethanol followed by cleavage after the His_6_-SUMO tag with heterologously expressed His_6_-tagged ULP1 protease, a kind gift from Birthe Kragelund. Ni^2+^-NTA resin was used to bind the protease and impurities bound to the Ni^2+^ NTA resin in the first affinity purification, and ALBA1 was collected in the flowthrough. ALBA1 was further purified on a HiLoad Superdex™ 200 10/300 GL prep grade column (GE Healtcare) connected to an HPLC ÄKTA Purifier system (GE Healthcare). Eluates were monitored at A_280_, and purity assessed by SDS-page analysis.

### Development of ALBA1 and ALBA4 antibodies

The anti-ALBA1, anti-ALBA4 antibodies were affinity-purified by Eurogentec from serum collected from rabbits immunized with recombinant ALBA1 protein or a 1:1 mix of the KLH-coupled ALBA4 peptides H-CGFNNRSDGPPVQAAA-OH and H- CNGPPNEYDAPQDGGY-NH_2_ (Eurogentech). The ALBA4 peptides were synthesized by Schafer-N Aps, Copenhagen, Denmark.

### Protein alignment and logo representation

Protein sequence fragments spanning the region from the N-terminal end of N8 to the C-terminal part of the YTH domain were aligned using ClustalW^69^. The high conservation of the YTH domains facilitated the definition of a common point of reference (the N-terminus of the YTH domain) for all protein sequences. Logo representations of the of the IDR parts of the alignment (ending in the common reference point) were made using Weblogo^68^. For alignment of ECT2 paralogs, Arabidopsis ECT1-ECT11 protein sequences were used. For alignment of ECT2 orthologs, we used the proteins with the highest fraction of sequence identity to Arabidopsis ECT2 (as determined by BlastP) from the following land plant species: *Marchantia polymorpha*, *Physcomitrella patens*, *Selaginella moellendorfii*, *Ceratopteris richardtii*, *Amborella trichopoda*, *Oryza sativa*, *Arabidopsis thaliana*.

### Structural modeling using AlphaFold3

The structural model of the ECT2-(ALBA4)_2_-RNA complex was generated by AlphaFold3^45^ using default settings and the following sequence input: 1 molecule of ECT2 (gene model AT3G13460.1, amino acid residues 373-616), 2 molecules of ALBA5 (gene model AT1G20220.1, amino acid residues 18-114), 1 molecule of RNA (5’-AAA[m^6^A]CUUCUG- 3’).

### ALBA4-GFP iCLIP experiments and library preparation

iCLIP experiments were carried out based on the method previously employed for Arabidopsis GRP7-GFP^50^ and the optimized iCLIP2 protocol^51,52^. Briefly, 7-day-old seedlings expressing ALBA4-GFP or GFP alone grown at 20°C in LD (16h light, 8h dark) were crosslinked with 254 nm UV light at 2000 mJ/cm^2^, snap frozen and ground into a fine powder in liquid nitrogen, and homogenized in iCLIP lysis buffer (50 mM Tris-HCl pH 7.5, 150 mM NaCl, 4 mM MgCl_2_, 5 mM DTT, 1% SDS, 0.25% sodium deoxycholate, 0.25% Igepal) supplemented with Roche Complete Protease Inhibitor cocktail (1 tablet/50 mL). The lysate was cleared by centrifugation and filtration (0.45 μm pore) of the supernatant. After pre-clearing with 200 μL of sepharose beads for 1h at 4°C, RNP-complexes were immunopurified with GFP-Trap beads (ChromoTek) for 4 hr at 4°C under constant rotation. We used 50 μL of beads for 3 g of tissue in 5 mL of iCLIP lysis buffer for every replicate. After washing four times with iCLIP wash buffer (2 M urea, 50 mM Tris-HCl pH 7.5, 500 mM NaCl, 4 mM MgCl_2_, 2 mM DTT, 1% SDS, 0.5% sodium deoxycholate, 0.5% Igepal, supplemented with Roche Complete Protease Inhibitor cocktail (1 tablet/50 mL)), and twice with PNK wash buffer (20 mM Tris-HCl, pH 7.4, 10 mM MgCl_2_, 0.2% Tween 20), RNP complexes attached to the beads were subjected to treatment with DNase (Turbo DNase [Ambion], 4 U/100 μL) and optionally RNase I (Ambion, 1 U/mL) at 37°C for 10 min. Subsequently, RNA 3′-ends were dephosphorylated (PNK [ThermoFisher] in buffer containing 350 mM Tris-HCl pH 6.5, 50 mM MgCl_2_, 25 mM DTT) for 20 min at 37°C, followed by one wash with PNK wash buffer, one wash with high-salt buffer (50 mM Tris-HCl pH 7.4, 1 M NaCl, 1 mM EDTA, 1% Igepal, 0.1% SDS, 0.5 % sodium deoxycholate) and two more washes with PNK wash buffer. The L3 linker was then ligated to the 3′ RNA ends (with NEB HC RNA Ligase in ligation buffer (200 mM Tris-HCl pH 7.8, 40 mM MgCl_2_, 40 mM DTT with RiboLock and PEG8000) at 16°C and 1250 rpm for >16h.

Samples were then washed twice in high-salt buffer and once in PNK wash buffer before the RNA was radioactively labeled at the 5′-end by PNK-mediated phosphorylation using L-^32^P- ATP (20 min at 37°C). The labeled RNP complexes were subjected to SDS-PAGE (4-12% NuPAGE Bis-Tris gel with 1x MOPS buffer) and blotting on a nitrocellulose membrane (Protran BA-85). Pieces of membrane containing a size range of RNA species bound to the protein (a smear above the expected molecular weight localized by autoradiography) were excised and subjected to proteolysis (200 μg of Proteinase K [Roche] in 200 μL of PK buffer [100 mM Tris-HCl pH 7.4, 50 mM NaCl, 10 mM EDTA] for 20 min at 37°C) to release RNA bound to small peptides. The RNA was then purified using phenol-chloroform (pH 7.0) and ethanol precipitation and used to prepare sequencing libraries following the iCLIP2 protocol^51^: reverse transcription with SSIII (Invitrogen) and an RT oligo complementary to the L3 liker followed by RNA hydrolysis and cDNA clean-up with MyONE Silane beads (Thermo Fisher). A second adapter was then ligated to the 3’OH of the cDNAs (with NEB HC RNA Ligase in NEB ligation buffer plus 5% DMSO, 1 mM ATP and 22.5% PEG8000) at 20°C and 1250 rpm overnight. The adapter contains a bipartite unique molecular identifier (UMI) and an experimental barcode, allowing for PCR duplicate removal and sample multiplexing, respectively. After another MyONE Silane clean-up, the cDNA library is pre-amplified in a first PCR (6 cycles) followed by size selection with ProNex beads (Promega) to remove short cDNAs and primer dimers. The cDNA library is then amplified in a second PCR followed by a second ProNex size selection to remove PCR primers and finally prepare the cDNA library for sequencing. The 2^nd^ PCR was carried out with 10 μL of cDNA and 8 cycles for each replicate. Samples were multiplexed and sequenced in the NextSeq sequencer (NextSeq® 500/550 Mid Output Kit v2 (150 cycles)) at the Genomics Core Facility at IMB (Mainz, Germany).

### ALBA4-GFP iCLIP analysis

All reads from iCLIP experiments were quality checked after multiple processing steps with FastQC (0.11.9). The distribution of read counts assigned to sample barcodes was computed using awk (GNU awk 5.0.1). Reads were demultiplexed, sequencing adapters removed from 3’ ends and subsequently quality- as well as length-trimmed (--min-read-length 15 -q WIN -qf sanger –min-read-length 15) with Flexbar (3.5.0) while keeping the random UMI parts in the read id field (--umi-tags). A genome index was created using STAR (2.7.3a) using the *Arabidopsis thaliana* genome version TAIR10. The genome annotation from Araport (version 11) was specified to mark the location splice junctions. Quality trimmed reads were then mapped using STAR and the created genome index, allowing only softclipping of 3’ ends (--alignEndsType Extend5pOfRead1) to preserve the position of the crosslinked nucleotide. PCR duplicates were removed using umi_tools (1.0.1) by considering the UMI tag in the read id field and the mapping coordinates. The uniquely mapped and deduplicated reads from each ALBA4-GFP and GFP replicate were merged together using samtools (1.14) and peak called with PureCLIP (1.3.1) in standard mode (-bc 0) to identify short and defined peak coordinates. In order to learn the HMM parameters only the first two chromosomes were specified (-iv ’Chr1;Chr2’) and the precision to store probabilities was set to long double (-ld). Clusters of directly adjacent called peaks were merged and reduced to the position with the highest reported PureCLIP score (1-nt resolution). Binding sites were defined as called peaks, extended by 4 nt (- 4…0…+4) in both directions with bedtools (2.27.1). Sites which reported crosslinks in only 1 out of 9 position were removed as they are considered artifacts. To confirm that the binding sites are supported by at least 2 replicates and a sufficient number of reads (reproducible binding sites), the coordinates of binding sites were overlapped with crosslink positions from every replicate (ALBA4-GFP and GFP independently). The distribution of crosslinks per binding sites was used to determine a reproducibility threshold. After defining a distribution quantile of 30% as the minimal filtering threshold, only binding sites above this threshold in at least 2 out of 3 replicates were kept. Due to the low amount of uniquely mapped reads the GFP control was not tested for reproducibility. Reproducible binding sites of ALBA4-GFP overlapping with binding sites from the GFP control were removed using bedtools and reported in browser extensible data (BED) format. Targets of ALBA4-GFP were defined as transcripts overlapping reproducible binding sites. Only the locations of representative gene models from Araport (version 11) were considered. For visual inspection data tracks were generated from uniquely mapped ALBA4-GFP and GFP only reads using bedtools.

### Sample preparation for TRIBE and HyperTRIBE

RNA extraction and library preparation was performed as previously described^28^. Total RNA was extracted from manually dissected root tips for ALBA4-FLAG-ADAR and apices (removing cotyledons) for ALBA2-FLAG-ADAR and ECT2-FLAG-ADAR of five independent lines (10-day-old T2 seedlings) with each of the lines being used as biological replicate.

### TRIBE/HyperTRIBE analyses for ALBA2 and ALBA4 vs. free ADAR controls

For all TRIBE/HyperTRIBE experiments, reads were mapped to the TAIR10 genome using STAR^77^ (version 2.7.11) and transcripts quantified using Salmon^78^ based on the Araport11 transcriptome^79^ augmented with the DNA sequence for the ADAR clone. The hyperTRIBER pipeline ^57^ was employed in order to quantify all positions with at least one mismatch to the genome, filter candidate positions by mutation type (A-to-G or T-to-C for forward or reverse strands, respectively) and replicate agreement, and formally test these candidates using a generalised linear model based approach for assessing difference in editing proportions between free ADAR control samples vs. fusion samples, retaining positions with a log_2_FC>1, an adjusted p-value <0.01 and a minimum editing proportion of 0.01. All sets were further annotated using the hyperTRIBER pipeline based on Araport11 gene annotations and prioritising highly expressed transcripts in the control lines in the case of positions overlapping multiple transcripts.

#### HyperTRIBE analysis for ECT2 on *alba1245* background and ALBA2 on *gte234* background

Unequal levels of *ECT2-FLAG-ADAR* or *ALBA2-FLAG-ADAR* expression between different genetic backgrounds in the same HyperTRIBE experiment could result in misinterpretation of results due to biased ADAR-driven editing patterns. This was supported by inspection of the initial results from the hyperTRIBER pipeline^57^ when comparing *ECT2-FLAG-ADAR-*expressing plants in the Col-0 vs *alba1245* backgrounds.

This preliminary analysis showed stronger editing in the direction of the samples with higher average *ADAR* expression, supported by western blots. To investigate further, we first re-ran the pipeline on only four lines (two per genetic background), selected such that the average number of reads mapping to *ADAR* was approximately equal between the two genetic backgrounds. Compared to the naïve analysis of all five lines per genotype, the significantly differently edited sites were visually less biased in the direction high *ADAR* expression, indicating that unequal *ADAR* expression leads to spurious results if left uncorrected. Furthermore, we observed a pattern whereby sites on lowly expressed genes tended to exhibit a larger editing proportion. To robustly account for differences in *ADAR* expression as measured by mRNA-seq read counts, we formulated a Bayesian hierarchical model as follows. First, we split the samples into three groups according to the (EXPR_BIN). Let *Y_ijkc_* denote the observed count of base *G* at the *i*-th position, with the *j*-expression of the ADAR clone (ADAR_BIN) and binned expression levels into 5 groups th level of ADAR_BIN, the *k*-th level of EXPR_BIN, and under condition *c*. *Y_ijkc_* is assumed to follow a Binomial distribution *Y_ijkc_* ∼ Binomial(*n_ijkc_*, *p_ijkc_*) where *n_ijkc_* represents the number of trials for each combination of position, ADAR_BIN level, EXPR_BIN level, and condition, and *p_ijkc_* is the probability of observing base *G*. Then the logit of *p_ijkc_* is modelled as 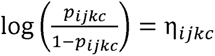 where the linear predictor *n_ijkc_* is given by:

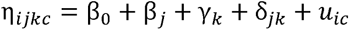

where β_0_ is the intercept, β*_j_* is the effect of ADAR bin *j*, γ*_k_* is the effect of expression bin *k*, *δ_jk_* is the corresponding ADAR expression interaction and 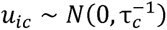 is a position-specific random effect with condition-specific precision parameter τ*_c_*. The model was fit using the Integrated Nested Latent Laplace (INLA) framework.

Let *u_iA_* and *u_iB_* denote the random effects for position *i* under conditions A and B, then the linear combination is LC*_i_* = *u_iA_* − *u_iB_* was computed from the posterior distribution of the fitted model. The mean 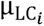 and standard deviation 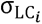 of samplings from the fitted posterior were used to generate Z-scores 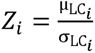 which were converted into p-values and subsequently adjusted to a false discovery rate (FDR). Importantly, the list of significant genes from this analysis strongly overlapped with the smaller list of genes from the 2-sample analysis described above (Supplementary Figure S11 H+I, compared to overlap with randomly sampled positions).

Finally, position-specific corrected editing proportions from the fitted model were further estimated by assuming ADAR to be exactly to the center bin and used for producing scatter plots for all tested positions.

### Definitions of strict and permissive gene sets

Strict sets: ALBA4, intersection of iCLIP (strong) and ALBA4 TRIBE associated gene sets. ECT2, intersection between ECT2/3 HyperTRIBE and ECT2 iCLIP (110 KDa) target sets^44^. Permissive sets: union instead of intersection between above sets for ALBA4 and ECT2, respectively.

### Venn diagrams and significance of overlaps

Venn diagrams were generated using custom code and the R-package eulerr (https://CRAN.R-project.org/package=eulerr)^80,81^. To assess the significance of overlaps between two sets of genes, a random set of genes of size equal to the number of genes in the first set was selected. To avoid expression bias—due to random genes being on average more lowly expressed than the sets of interest—the expression distribution of the random set was matched to that of the first set. We calculated the number of genes in the first set overlapping with the second set, as well as the number of genes for each of 1000 random samples overlapping with the second set. The *p*-value was calculated as: 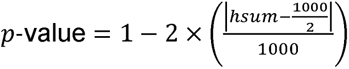 where hsum is the number of cases where the number of genes in the random set overlapped more with the second set. In cases where there were zero instances where the random set had a better overlap with the second set than the first set, the *p*-value was set to "<0.001", indicating a high significance of overlap. This procedure was carried out using a custom script, which also returned a single random set of (expression distribution matched) genes. This random set was used in the Venn diagrams to provide a visual indication of the expected overlap by chance.

In order to check for possible false positives in genes with fewer than 2-5 iCLIP sites, we overlapped the set with the ALBA4 HyperTRIBE data and looked for the percentage of support. We noted that genes with only a single, low quality iCLIP site tended to be supported by ALBA4 HyperTRIBE to a similar level as random sets of expressed genes, providing justification for considering the more robust set for subsequent analyses.

### Metagene plots

Metagene plots showing enrichment of features in 5’-UTR, CDS and 3’-UTR corrected for the size of the annotated region were generated using a strategy similar to what we previously reported^28^.

### Single cell co-expression analysis of ECT2

We first obtained single-cell mRNA-seq root tip data^46,47^. To avoid bias due differences in UMI count between ECT2 expressing (ECT2+) cells and non-expressing (ECT2-) cells, each ECT2+ cell was matched with an ECT2- cell of similar UMI count. For each gene G expressed within the range of 20-80% of the resulting total cells, counts of G+ and G- cells for each of the ECT2+ and ECT2- sets were used to perform a fisher’s exact test, whereby a high odds ratio represents a high corresponding between ECT2 and the tested G, indicative of co-expression.

### Motif analysis

We first considered the set of motifs previously defined on the basis of ECT2 iCLIP data^28^. Background sites for m^6^A (nanopore-derived [54]), ALBA4 iCLIP and ECT2 iCLIP were generated following a similar strategy to what we previously reported^28^, ensuring that the distribution of site locations across gene features were identical for both the true set and the background set. We subsequently removed background sites which by chance overlapped with sites from the true sets (within 100 bp). For both the true sets and background sites, we calculated the number of motifs present per 1000 sites (a normalization allowing for comparability across different sets), for each position up to 100 bp from the site.

### Curation of m^6^A site set

We first collected m^6^A sites for *A. thaliana* from multiple published sources^54,82^. As nanopore-derived sites are not subjected to UV-bias, we trained a neural network to differentiate between the 20,858 m^6^A sites identified by nanopore^54^ and a corresponding set of 20,715 location-matched negative sites (Figure S6A). The neural network used as input extracted sequence +/-100 bp regions around all positions (R-packages BSgenome^83^ and AThaliana), which was converted from FASTA to one-hot encoded format^84^. As output, the network predicted the presence or absence of the m6A at the center point of the input sequence. The network was based on 4 blocks of: 1D convolutional layer with relu activation, batch normalization and max pooling of size 2. The output of these four blocks was flattened, run through a fully connected layer and then passed into a fully connected output layer of output size 1 with a sigmoid activation function. The model was trained specifying the binary cross-entropy loss function using Keras with a Tensorflow back-end^85^ specifying binary cross entropy loss function. The model showed excellent performance, with AUC ranging 0.85-0.92 over the five folds. This model enabled us to fine-adjust sites from other sets by systematically shifting their positions and selecting those with the highest probability (Figure S6B, Methods). Consequently, we augmented the smaller set of nanopore-derived positions with a broader set exhibiting properties highly consistent with nanopore-identified sites (Figure S6B-E) Notably, approx. 90,000 miCLIP-derived positions not only shifted to locations similar to nearby nanopore-defined sites, but also consolidated into fewer positions, indicating that many miCLIP-identified sites represent imprecise locations. Overall, our augmentation strategy yielded a compendium of 41,883 m^6^A sites in *A. thaliana*.

### Convolutional neural network based de-novo motif detection

We annotated each of 41,883 m^6^A sites according to overlap of either ECT2 iCLIP or ALBA iCLIP sites within 100 bp. For each m6A site, 300 bp of sequence was extracted either side, creating a 601 bp long sequences, which were embedded using one-hot encoding and passed as input in a convolutional neural network with two outputs - presence or absence of ECT2 iCLIP and presence or absence of ALBA iCLIP. The network architecture consisted of five blocks of: a 1D convolutional layers with RELU activation, 0.2 drop-out layer, a batch normalisation layer and a max-pooling layer with pool size 2. Each convolutional layer had 64 filters, with a kernal size of 8 in the first layer and 6 thereafter. The output was then flattened into one dimension and passed through a separate a connected layer of kernel 32 for each output, which was specified as a fully connected layer of size 1 using a sigmoid activation function. The network was trained using Keras with a Tensorflow back-end^85^, specifying the binary cross entropy loss function for each output.

### 5-fold cross-validation strategy for machine learning models

Sites were split into 5 sets of similar size. Since there are often multiple m6A sites on a single gene, and these sites often fall within overlapping windows, we separated training and test sets such that no gene was present in both sets. Each testing set consisted of one of the 5 sets, and the training set the remaining sets combined. All predictions used in subsequent analyses were based only on sets held out of the training process.

### Modelling of RBP-specific motifs

After fitting each fold weights for the 32 learned convolutional filters of length 8 from the initial layer (that is, the layer connecting to the input sequence) were extracted, resulting in a total of 160 filters. For each of these filters individually, we scanned through all sequences from the training set, selected the top 5000 high-scoring positions and used the resulting nucleotide frequencies at each of the 8 positions to derive a position weight matrix. These position weight matrices were then allocated a consensus name using the R package universalmotif ^86^.

For each motif, m^6^A-centered sequences were classified as containing or not-containing the given motif within 150 bp of the methylation site. In order to detect RBP-specific binding motifs, a generalized linear model (glm) assuming a binomial-distributed response was used to predict motif presence as the dependent variable, where the two predictors in the model were the probability of ECT2 binding from the neural network and the probability of ALBA4 binding from the neural network. In this way, the coefficient for ECT2 binding is interpreted as the strength of correspondence with that motif whilst controlling for binding of ALBA4, and vice-versa. Z-scores for each of the two proteins for all motifs were then extracted from the model and plotted as enrichment scores.

## ACKNOWLEDGEMENTS

We thank Kristina Neudorf, Lena Bjørn Johansson, Daniel Tobias Kyndesen Lahti, Ida Thorøe Michler, Magnus von Holstein-Rathlou, and Jakub Najbar for their valuable technical assistance and Theo Bölsterli and René Hvidberg Petersen and their teams for plant care. Christian Poulsen is thanked for help with AlphaFold3 modeling. We acknowledge Erwin Schoof and the proteomics platform at Denmark’s Technical University for their expertise in running protein identification through liquid chromatography-mass spectrometry. We also thank Mandy Rettel and Frank Stein from the EMBL Proteomics Core facility for IP-MS analysis of ALBA4 IPs. Carlotta Porcelli is thanked for advice on analysis of mass spectrometry data. Support by the IMB Genomics Core Facility and the use of its NextSeq500 (funded by the Deutsche Forschungsgemeinschaft (DFG, German Research Foundation) – INST 247/870-1 FUGG) is gratefully acknowledged. This research was supported by a Hallas-Møller Ascending Investigator Fellowship grant from the Novo Nordisk Foundation (NNF19OC0054973), a Consolidator Grant from the European Research Council (PATHORISC, ERC-2016-CoG 726417), a Research Infrastructure Grant from Carlsberg Fondet (CF20-0659), and an Instrument Grant from Brdr Hartmann Fonden (A35879), all awarded to PB, and by grants STA653/13 and STA653/14 from Deutsche Forschungsgemeinschaft to DS.

## AUTHOR CONTRIBUTIONS

M.R. carried out IP-MS and iCLIP experiments with ALBA4-GFP and noticed that ECT proteins were of particular interest for follow-up studies, conducted crosslinking-PNK labeling experiments, and characterized *alba* mutant phenotypes. M.D.T. made and characterized the *ect2-5* in-frame deletion mutant, made transgenic lines expressing ECT2-mCherry mutants and used them for IP-MS and IP-western blot analyses, noticed that ALBA proteins were of particular interest for follow-up study, made ALBA1/4 antibodies, conducted TRIBE and HyperTRIBE experiments, and made and analysed ALBA1-TFP and ALBA4-Venus transgenic lines. S.R. designed and carried out all computational analyses except iCLIP peak calling, with most objectives determined in discussion with L.A-H, M.R., M.D.T. and P.B. M.L. called ALBA4 iCLIP peaks. T.K. provided guidance on iCLIP optimization for ALBA4-GFP. L.A-H designed initial steps of ECT2 N8 characterization with P.B., supervised M.D.T.’s work towards these goals, and participated in decisions on project directions and data presentation. N.W. constructed composite *alba* mutants, and constructed and analyzed ALBA-GFP-expressing transgenic lines. T.M. supervised work on construction of *alba* mutants, ALBA-GFP transgenic lines, and confocal microscopy of ALBA-GFP lines, D.S. supervised ALBA4 IP-MS and iCLIP experiments, P.B. designed, supervised and coordinated the project, and wrote the manuscript together with S.R., M.R. and M.D.T. All authors contributed improvements on the first manuscript draft.

## DATA AVAILABILITY

All sequencing data have been deposited in the European Nucleotide Archive under accession code PRJEB71752. The mass spectrometry proteomics data have been deposited to the ProteomeXchange Consortium via the PRIDE^87^ partner repository with the dataset identifier PXD052232.

### Reviewer access

**Password:** XAPWwwxp

Code used for data analysis is available at Github: https://github.com/sarah-ku/ALBA_YTH_arabidopsis

## REFERENCES

1. Balacco, D.L., and Soller, M. (2019). The m6A Writer: Rise of a Machine for Growing Tasks. Biochemistry 58, 363–378. 10.1021/acs.biochem.8b01166.

2. Zhong, S., Li, H., Bodi, Z., Button, J., Vespa, L., Herzog, M., and Fray, R.G. (2008). MTA is an Arabidopsis messenger RNA adenosine methylase and interacts with a homolog of a sex-specific splicing factor. The Plant cell 20, 1278–1288. 10.1105/tpc.108.058883.

3. Geula, S., Moshitch-Moshkovitz, S., Dominissini, D., Mansour, A.A., Kol, N., Salmon-Divon, M., Hershkovitz, V., Peer, E., Mor, N., Manor, Y.S., et al. (2015). m6A mRNA methylation facilitates resolution of naive pluripotency toward differentiation. Science 347, 1002–1006. 10.1126/science.1261417.

4. Clancy, M.J., Shambaugh, M.E., Timpte, C.S., and Bokar, J.A. (2002). Induction of sporulation in Saccharomyces cerevisiae leads to the formation of N6-methyladenosine in mRNA: a potential mechanism for the activity of the IME4 gene. Nucleic acids research 30, 4509–4518.

5. Lence, T., Akhtar, J., Bayer, M., Schmid, K., Spindler, L., Ho, C.H., Kreim, N., Andrade-Navarro, M.A., Poeck, B., Helm, M., and Roignant, J.-Y. (2016). m6A modulates neuronal functions and sex determination in Drosophila. Nature 540, 242–247. 10.1038/nature20568.

6. Haussmann, I.U., Bodi, Z., Sanchez-Moran, E., Mongan, N.P., Archer, N., Fray, R.G., and Soller, M. (2016). m6A potentiates Sxl alternative pre-mRNA splicing for robust Drosophila sex determination. Nature 540, 301–304. 10.1038/nature20577.

7. Arribas-Hernández, L., Bressendorff, S., Hansen, M.H., Poulsen, C., Erdmann, S., and Brodersen, P. (2018). An m^6^A-YTH Module Controls Developmental Timing and Morphogenesis in Arabidopsis. The Plant cell 30, 952–967.

8. Ivanova, I., Much, C., Di Giacomo, M., Azzi, C., Morgan, M., Moreira, P.N., Monahan, J., Carrieri, C., Enright, A.J., and O’Carroll, D. (2017). The RNA m(6)A Reader YTHDF2 Is Essential for the Post-transcriptional Regulation of the Maternal Transcriptome and Oocyte Competence. Molecular cell 67, 1059–1067 e1054. 10.1016/j.molcel.2017.08.003.

9. Lasman, L., Krupalnik, V., Viukov, S., Mor, N., Aguilera-Castrejon, A., Schneir, D., Bayerl, J., Mizrahi, O., Peles, S., Tawil, S., et al. (2020). Context-dependent functional compensation between Ythdf m(6)A reader proteins. Genes & development 34, 1373–1391. 10.1101/gad.340695.120.

10. Kontur, C., Jeong, M., Cifuentes, D., and Giraldez, A.J. (2020). Ythdf m6A Readers Function Redundantly during Zebrafish Development. Cell reports 33. 10.1016/j.celrep.2020.108598.

11. Patil, D.P., Pickering, B.F., and Jaffrey, S.R. (2018). Reading m(6)A in the Transcriptome: m(6)A-Binding Proteins. Trends Cell Biol 28, 113–127. 10.1016/j.tcb.2017.10.001.

12. Fray, R.G., and Simpson, G.G. (2015). The Arabidopsis epitranscriptome. Current Opinion in Plant Biology 27, 17–21. 10.1016/j.pbi.2015.05.015.

13. Scutenaire, J., Deragon, J.-M., Jean, V., Benhamed, M., Raynaud, C., Favory, J.-J., Merret, R., and Bousquet-Antonelli, C. (2018). The YTH Domain Protein ECT2 Is an m^6^A Reader Required for Normal Trichome Branching in Arabidopsis. The Plant cell 30, 986.

14. Ok, S.H., Jeong, H.J., Bae, J.M., Shin, J.S., Luan, S., and Kim, K.N. (2005). Novel CIPK1-associated proteins in Arabidopsis contain an evolutionarily conserved C-terminal region that mediates nuclear localization. Plant physiology 139, 138–150. 10.1104/pp.105.065649.

15. Arribas-Hernandez, L., Simonini, S., Hansen, M.H., Paredes, E.B., Bressendorff, S., Dong, Y., Ostergaard, L., and Brodersen, P. (2020). Recurrent requirement for the m(6)A-ECT2/ECT3/ECT4 axis in the control of cell proliferation during plant organogenesis. Development 147. 10.1242/dev.189134.

16. Yin, S., Ao, Q., Qiu, T., Tan, C., Tu, Y., Kuang, T., and Yang, Y. (2022). Tomato SlYTH1 encoding a putative RNA m6A reader affects plant growth and fruit shape. Plant Science 323, 111417. 10.1016/j.plantsci.2022.111417.

17. Ma, W., Cui, S., Lu, Z., Yan, X., Cai, L., Lu, Y., Cai, K., Zhou, H., Ma, R., Zhou, S., and Wang, X. (2022). YTH Domain Proteins Play an Essential Role in Rice Growth and Stress Response. Plants 11. 10.3390/plants11172206.

18. Su, D., Yang, L., Shi, X., Ma, X., Zhou, X., Hedges, S.B., and Zhong, B. (2021). Large-Scale Phylogenomic Analyses Reveal the Monophyly of Bryophytes and Neoproterozoic Origin of Land Plants. Molecular Biology and Evolution 38, 3332–3344. 10.1093/molbev/msab106.

19. Magallón, S., Hilu, K.W., and Quandt, D. (2013). Land plant evolutionary timeline: Gene effects are secondary to fossil constraints in relaxed clock estimation of age and substitution rates. American Journal of Botany 100, 556–573. 10.3732/ajb.1200416.

20. Flores-Téllez, D., Tankmar, M.D., von Bülow, S., Chen, J., Lindorff-Larsen, K., Brodersen, P., and Arribas-Hernández, L. (2023). Insights into the conservation and diversification of the molecular functions of YTHDF proteins. PLoS genetics 19, e1010980. 10.1371/journal.pgen.1010980.

21. Tankmar, M.D., Reichel, M., Arribas-Hernández, L., and Brodersen, P. (2023). A YTHDF-PABP interaction is required for m6A-mediated organogenesis in plants. EMBO reports 24, e57741. 10.15252/embr.202357741.

22. Song, P., Wei, L., Chen, Z., Cai, Z., Lu, Q., Wang, C., Tian, E., and Jia, G. (2023). m(6)A readers ECT2/ECT3/ECT4 enhance mRNA stability through direct recruitment of the poly(A) binding proteins in Arabidopsis. Genome biology 24, 103. 10.1186/s13059-023-02947-4.

23. Wiedner, H.J., and Giudice, J. (2021). It’s not just a phase: function and characteristics of RNA-binding proteins in phase separation. Nature structural & molecular biology 28, 465–473. 10.1038/s41594-021-00601-w.

24. Lee, K.P., Liu, K., Kim, E.Y., Medina-Puche, L., Dong, H., Di, M., Singh, R.M., Li, M., Qi, S., Meng, Z., et al. (2023). The m6A reader ECT1 drives mRNA sequestration to dampen salicylic acid–dependent stress responses in Arabidopsis. The Plant cell 36, 746–763. 10.1093/plcell/koad300.

25. Wu, X., Su, T., Zhang, S., Zhang, Y., Wong, C.E., Ma, J., Shao, Y., Hua, C., Shen, L., and Yu, H. (2024). N6-methyladenosine-mediated feedback regulation of abscisic acid perception via phase-separated ECT8 condensates in Arabidopsis. Nature Plants 10, 469–482. 10.1038/s41477-024-01638-7.

26. Stowell, J.A.W., Wagstaff, J.L., Hill, C.H., Yu, M., McLaughlin, S.H., Freund, S.M.V., and Passmore, L.A. (2018). A low-complexity region in the YTH domain protein Mmi1 enhances RNA binding. Journal of Biological Chemistry 293, 9210–9222. 10.1074/jbc.RA118.002291.

27. Chong, P.A., Vernon, R.M., and Forman-Kay, J.D. (2018). RGG/RG Motif Regions in RNA Binding and Phase Separation. Journal of Molecular Biology 430, 4650–4665. 10.1016/j.jmb.2018.06.014.

28. Arribas-Hernández, L., Rennie, S., Köster, T., Porcelli, C., Lewinski, M., Staiger, D., Andersson, R., and Brodersen, P. (2021). Principles of mRNA targeting via the Arabidopsis m6A-binding protein ECT2. eLife 10, e72375. 10.7554/eLife.72375.

29. Holehouse, A.S., and Kragelund, B.B. (2024). The molecular basis for cellular function of intrinsically disordered protein regions. Nature Reviews Molecular Cell Biology 25, 187–211. 10.1038/s41580-023-00673-0.

30. Reichel, M., Liao, Y., Rettel, M., Ragan, C., Evers, M., Alleaume, A.M., Horos, R., Hentze, M.W., Preiss, T., and Millar, A.A. (2016). In Planta Determination of the mRNA-Binding Proteome of Arabidopsis Etiolated Seedlings. The Plant cell 28, 2435–2452. 10.1105/tpc.16.00562.

31. Marondedze, C., Thomas, L., Serrano, N.L., Lilley, K.S., and Gehring, C. (2016). The RNA-binding protein repertoire of Arabidopsis thaliana. Scientific Reports 6, 29766. 10.1038/srep29766.

32. Aravind, L., Iyer, L.M., and Anantharaman, V. (2003). The two faces of Alba: the evolutionary connection between proteins participating in chromatin structure and RNA metabolism. Genome biology 4, R64. 10.1186/gb-2003-4-10-r64.

33. Forterre, P., Confalonieri, F., and Knapp, S. (1999). Identification of the gene encoding archeal-specific DNA-binding proteins of the Sac10b family. Molecular Microbiology 32, 669–670. 10.1046/j.1365-2958.1999.01366.x.

34. Xue, H., Guo, R., Wen, A., Liu, D., and Huang, L. (2000). An Abundant DNA Binding Protein from the Hyperthermophilic Archaeon *Sulfolobus shibatae* Affects DNA Supercoiling in a Temperature-Dependent Fashion. Journal of bacteriology 182, 3929–3933. doi:10.1128/jb.182.14.3929-3933.2000.

35. Bell, S.D., Botting, C.H., Wardleworth, B.N., Jackson, S.P., and White, M.F. (2002). The Interaction of Alba, a Conserved Archaeal Chromatin Protein, with Sir2 and Its Regulation by Acetylation. Science 296, 148–151. doi:10.1126/science.1070506.

36. Wardleworth, B.N., Russell, R.J.M., Bell, S.D., Taylor, G.L., and White, M.F. (2002). Structure of Alba: an archaeal chromatin protein modulated by acetylation. The EMBO journal 21, 4654–4662. 10.1093/emboj/cdf465.

37. Zhang, N., Guo, L., and Huang, L. (2020). The Sac10b homolog from Sulfolobus islandicus is an RNA chaperone. Nucleic acids research 48, 9273–9284. 10.1093/nar/gkaa656.

38. Goyal, M., Banerjee, C., Nag, S., and Bandyopadhyay, U. (2016). The Alba protein family: Structure and function. Biochimica et Biophysica Acta (BBA) - Proteins and Proteomics 1864, 570–583. 10.1016/j.bbapap.2016.02.015.

39. Mani, J., Güttinger, A., Schimanski, B., Heller, M., Acosta-Serrano, A., Pescher, P., Späth, G., and Roditi, I. (2011). Alba-Domain Proteins of Trypanosoma brucei Are Cytoplasmic RNA-Binding Proteins That Interact with the Translation Machinery. PloS one 6, e22463. 10.1371/journal.pone.0022463.

40. Bevkal, S., Naguleswaran, A., Rehmann, R., Kaiser, M., Heller, M., and Roditi, I. (2021). An Alba-domain protein required for proteome remodelling during trypanosome differentiation and host transition. PLOS Pathogens 17, e1009239. 10.1371/journal.ppat.1009239.

41. Honkanen, S., Jones, V.A.S., Morieri, G., Champion, C., Hetherington, A.J., Kelly, S., Proust, H., Saint-Marcoux, D., Prescott, H., and Dolan, L. (2016). The Mechanism Forming the Cell Surface of Tip-Growing Rooting Cells Is Conserved among Land Plants. Curr. Biol. 26, 3238–3244. 10.1016/j.cub.2016.09.062.

42. Magwanga, R.O., Kirungu, J.N., Lu, P., Cai, X., Xu, Y., Wang, X., Zhou, Z., Hou, Y., Agong, S.G., Wang, K., and Liu, F. (2019). Knockdown of ghAlba_4 and ghAlba_5 Proteins in Cotton Inhibits Root Growth and Increases Sensitivity to Drought and Salt Stresses. Frontiers in Plant Science 10. 10.3389/fpls.2019.01292.

43. Tong, J., Ren, Z., Sun, L., Zhou, S., Yuan, W., Hui, Y., Ci, D., Wang, W., Fan, L.-M., Wu, Z., and Qian, W. (2022). ALBA proteins confer thermotolerance through stabilizing HSF messenger RNAs in cytoplasmic granules. Nature Plants 8, 778–791. 10.1038/s41477-022-01175-1.

44. Arribas-Hernández, L., Rennie, S., Schon, M., Porcelli, C., Enugutti, B., Andersson, R., Nodine, M.D., and Brodersen, P. (2021). The YTHDF proteins ECT2 and ECT3 bind largely overlapping target sets and influence target mRNA abundance, not alternative polyadenylation. eLife 10, e72377. 10.7554/eLife.72377.

45. Abramson, J., Adler, J., Dunger, J., Evans, R., Green, T., Pritzel, A., Ronneberger, O., Willmore, L., Ballard, A.J., Bambrick, J., et al. (2024). Accurate structure prediction of biomolecular interactions with AlphaFoldL3. Nature. 10.1038/s41586-024-07487-w.

46. Shahan, R., Hsu, C.-W., Nolan, T.M., Cole, B.J., Taylor, I.W., Greenstreet, L., Zhang, S., Afanassiev, A., Vlot, A.H.C., Schiebinger, G., et al. (2022). A single-cell *Arabidopsis* root atlas reveals developmental trajectories in wild-type and cell identity mutants. Developmental cell 57, 543–560.e549. 10.1016/j.devcel.2022.01.008.

47. He, Z., Luo, Y., Zhou, X., Zhu, T., Lan, Y., and Chen, D. (2023). scPlantDB: a comprehensive database for exploring cell types and markers of plant cell atlases. Nucleic acids research 52, D1629–D1638. 10.1093/nar/gkad706.

48. McMahon, A.C., Rahman, R., Jin, H., Shen, J.L., Fieldsend, A., Luo, W., and Rosbash, M. (2016). TRIBE: Hijacking an RNA-Editing Enzyme to Identify Cell-Specific Targets of RNA-Binding Proteins. Cell 165, 742–753. 10.1016/j.cell.2016.03.007.

49. Konig, J., Zarnack, K., Rot, G., Curk, T., Kayikci, M., Zupan, B., Turner, D.J., Luscombe, N.M., and Ule, J. (2010). iCLIP reveals the function of hnRNP particles in splicing at individual nucleotide resolution. Nature structural & molecular biology 17, 909–915. 10.1038/nsmb.1838.

50. Meyer, K., Köster, T., Nolte, C., Weinholdt, C., Lewinski, M., Grosse, I., and Staiger, D. (2017). Adaptation of iCLIP to plants determines the binding landscape of the clock-regulated RNA-binding protein AtGRP7. Genome biology 18, 204. 10.1186/s13059-017-1332-x.

51. Buchbender, A., Mutter, H., Sutandy, F.X.R., Körtel, N., Hänel, H., Busch, A., Ebersberger, S., and König, J. (2020). Improved library preparation with the new iCLIP2 protocol. Methods 178, 33–48. 10.1016/j.ymeth.2019.10.003.

52. Lewinski, M., Brüggemann, M., Köster, T., Reichel, M., Bergelt, T., Meyer, K., König, J., Zarnack, K., and Staiger, D. (2024). Mapping protein–RNA binding in plants with individual-nucleotide-resolution UV cross-linking and immunoprecipitation (plant iCLIP2). Nature protocols 19, 1183–1234. 10.1038/s41596-023-00935-3.

53. Xu, W., Rahman, R., and Rosbash, M. (2018). Mechanistic implications of enhanced editing by a HyperTRIBE RNA-binding protein. Rna 24, 173–182. 10.1261/rna.064691.117.

54. Parker, M.T., Knop, K., Sherwood, A.V., Schurch, N.J., Mackinnon, K., Gould, P.D., Hall, A.J., Barton, G.J., and Simpson, G.G. (2020). Nanopore direct RNA sequencing maps the complexity of Arabidopsis mRNA processing and m(6)A modification. Elife 9, 49658. 10.7554/eLife.49658.

55. Hafner, M., Katsantoni, M., Köster, T., Marks, J., Mukherjee, J., Staiger, D., Ule, J., and Zavolan, M. (2021). CLIP and complementary methods. Nature Reviews Methods Primers 1, 20. 10.1038/s43586-021-00018-1.

56. Angelov, D., Boopathi, R., Lone, I.N., Menoni, H., Dimitrov, S., and Cadet, J. (2023). Capturing Protein–Nucleic Acid Interactions by High-Intensity Laser-Induced Covalent Cross-Linking†. Photochemistry and Photobiology 99, 296–312. 10.1111/php.13699.

57. Rennie, S., Magnusson, D.H., and Andersson, R. (2021). hyperTRIBER: a flexible R package for the analysis of differential RNA editing. bioRxiv, 2021.2010.2020.465108. 10.1101/2021.10.20.465108.

58. Bodi, Z., Zhong, S., Mehra, S., Song, J., Graham, N., Li, H., May, S., and Fray, R.G. (2012). Adenosine Methylation in Arabidopsis mRNA is Associated with the 3’ End and Reduced Levels Cause Developmental Defects. Front Plant Sci 3, 48. 10.3389/fpls.2012.00048.

59. Wei, L.-H., Song, P., Wang, Y., Lu, Z., Tang, Q., Yu, Q., Xiao, Y., Zhang, X., Duan, H.-C., and Jia, G. (2018). The m6A Reader ECT2 Controls Trichome Morphology by Affecting mRNA Stability in Arabidopsis. The Plant cell 30, 968.

60. Jumper, J., Evans, R., Pritzel, A., Green, T., Figurnov, M., Ronneberger, O., Tunyasuvunakool, K., Bates, R., Žídek, A., Potapenko, A., et al. (2021). Highly accurate protein structure prediction with AlphaFold. Nature 596, 583–589. 10.1038/s41586-021-03819-2.

61. Sikorski, V., Selberg, S., Lalowski, M., Karelson, M., and Kankuri, E. (2023). The structure and function of YTHDF epitranscriptomic m^6^A readers. Trends in Pharmacological Sciences 44, 335–353. 10.1016/j.tips.2023.03.004.

62. Wang, C., Zhu, Y., Bao, H., Jiang, Y., Xu, C., Wu, J., and Shi, Y. (2016). A novel RNA-binding mode of the YTH domain reveals the mechanism for recognition of determinant of selective removal by Mmi1. Nucleic acids research 44, 969–982. 10.1093/nar/gkv1382.

63. Huang, H., Weng, H., Sun, W., Qin, X., Shi, H., Wu, H., Zhao, B.S., Mesquita, A., Liu, C., Yuan, C.L., et al. (2018). Recognition of RNA N6-methyladenosine by IGF2BP proteins enhances mRNA stability and translation. Nature cell biology 20, 285–295. 10.1038/s41556-018-0045-z.

64. Hafner, M., Landthaler, M., Burger, L., Khorshid, M., Hausser, J., Berninger, P., Rothballer, A., Ascano, M., Jr., Jungkamp, A.-C., Munschauer, M., et al. (2010). Transcriptome-wide Identification of RNA-Binding Protein and MicroRNA Target Sites by PAR-CLIP. Cell 141, 129–141. 10.1016/j.cell.2010.03.009.

65. Zaccara, S., and Jaffrey, S.R. (2020). A Unified Model for the Function of YTHDF Proteins in Regulating m(6)A-Modified mRNA. Cell 181, 1582–1595 e1518. 10.1016/j.cell.2020.05.012.

66. Di Domenico, T., Walsh, I., Martin, A.J.M., and Tosatto, S.C.E. (2012). MobiDB: a comprehensive database of intrinsic protein disorder annotations. Bioinformatics 28, 2080–2081. 10.1093/bioinformatics/bts327.

67. Schneider, T.D., and Stephens, R.M. (1990). Sequence logos: a new way to display consensus sequences. Nucleic acids research 18, 6097–6100. 10.1093/nar/18.20.6097.

68. Crooks, G.E., Hon, G., Chandonia, J.-M., and Brenner, S.E. (2004). WebLogo: A Sequence Logo Generator. Genome Res. 14, 1188–1190. 10.1101/gr.849004.

69. Thompson, J.D., Higgins, D.G., and Gibson, T.J. (1994). CLUSTAL W: improving the sensitivity of progressive multiple sequence alignment through sequence weighting, position specific gap penalties and weight matrix choice. Nucleic acids research 22, 4673–4680.

70. Tsutsui, H., and Higashiyama, T. (2017). pKAMA-ITACHI Vectors for Highly Efficient CRISPR/Cas9-Mediated Gene Knockout in Arabidopsis thaliana. Plant and Cell Physiology 58, 46–56. 10.1093/pcp/pcw191.

71. Bitinaite, J., and Nichols, N.M. (2009). DNA Cloning and Engineering by Uracil Excision. Current Protocols in Molecular Biology 86, 3.21.21-23.21.16. 10.1002/0471142727.mb0321s86.

72. Nour-Eldin, H.H., Hansen, B.G., Norholm, M.H., Jensen, J.K., and Halkier, B.A. (2006). Advancing uracil-excision based cloning towards an ideal technique for cloning PCR fragments. Nucleic acids research 34, e122. 10.1093/nar/gkl635.

73. Curtis, M.D., and Grossniklaus, U. (2003). A Gateway Cloning Vector Set for High-Throughput Functional Analysis of Genes in Planta. Plant physiology 133, 462–469. 10.1104/pp.103.027979.

74. Earley, K.W., Haag, J.R., Pontes, O., Opper, K., Juehne, T., Song, K., and Pikaard, C.S. (2006). Gateway-compatible vectors for plant functional genomics and proteomics. The Plant Journal 45, 616–629. 10.1111/j.1365-313X.2005.02617.x.

75. Clough, S.J., and Bent, A.F. (1998). Floral dip: a simplified method for Agrobacterium-mediated transformation of Arabidopsis thaliana. Plant J 16, 735–743.

76. Speth, C., Toledo-Filho, L.A.A., and Laubinger, S. (2014). Immunoprecipitation-Based Analysis of Protein–Protein Interactions. In Plant Circadian Networks: Methods and Protocols, D. Staiger, ed. (Springer New York), pp. 175-185. 10.1007/978-1-4939-0700-7_11.

77. Dobin, A., Davis, C.A., Schlesinger, F., Drenkow, J., Zaleski, C., Jha, S., Batut, P., Chaisson, M., and Gingeras, T.R. (2013). STAR: ultrafast universal RNA-seq aligner. Bioinformatics 29, 15–21. 10.1093/bioinformatics/bts635.

78. Patro, R., Duggal, G., Love, M.I., Irizarry, R.A., and Kingsford, C. (2017). Salmon provides fast and bias-aware quantification of transcript expression. Nature Methods 14, 417–419. 10.1038/nmeth.4197.

79. Cheng, C.-Y., Krishnakumar, V., Chan, A.P., Thibaud-Nissen, F., Schobel, S., and Town, C.D. (2017). Araport11: a complete reannotation of the Arabidopsis thaliana reference genome. The Plant Journal 89, 789–804. 10.1111/tpj.13415.

80. Larsson, J., Gustafsson, P. (2018). A case study in fitting area-proportional euler diagrams with ellipses using eulerr. CEUR Workshop Proceedings - Proceedings of International Workshop on Set Visualization and Reasoning 2116, 84–91.

81. Larsson, J. (2022). eulerr: Area-Proportional Euler and Venn Diagrams with Ellipses. .

82. Tang, Y., Chen, K., Song, B., Ma, J., Wu, X., Xu, Q., Wei, Z., Su, J., Liu, G., Rong, R., et al. (2020). m6A-Atlas: a comprehensive knowledgebase for unraveling the N6-methyladenosine (m6A) epitranscriptome. Nucleic acids research 49, D134–D143. 10.1093/nar/gkaa692.

83. Pagès, H. (2024). BSgenome: Software infrastructure for efficient representation of full genomes and their SNPs. R package version 1.70.2. Bioconductor. 10.18129/B9.bioc.BSgenome.

84. Pedregosa, F., Varoquaux, G., Gramfort, A., Michel, V., Thirion, B., Grisel, O., Blondel, M., Prettenhofer, P., Weiss, R., Dubourg, V., et al. (2011). Scikit-learn: Machine Learning in Python. J. Mach. Learn. Res. 12, 2825–2830.

85. Abadi, M., Barham, P., Chen, J., Chen, Z., Davis, A., Dean, J., Devin, M., Ghemawat, S., Irving, G., Isard, M., et al. (2016). TensorFlow: a system for large-scale machine learning. Proceedings of the 12th USENIX conference on Operating Systems Design and Implementation. USENIX Association.

86. Tremblay, B.J. (2024). universalmotif: Import, Modify, and Export Motifs with R. 10.18129/B9.bioc.universalmotif.

87. Perez-Riverol, Y., Bai, J., Bandla, C., García-Seisdedos, D., Hewapathirana, S., Kamatchinathan, S., Kundu, Deepti J., Prakash, A., Frericks-Zipper, A., Eisenacher, M., et al. (2021). The PRIDE database resources in 2022: a hub for mass spectrometry-based proteomics evidences. Nucleic acids research 50, D543–D552. 10.1093/nar/gkab1038.

