## Supplemental Figures S1-S12 for "ALBA proteins facilitate cytoplasmic YTHDF-mediated reading of m^6^A in plants"

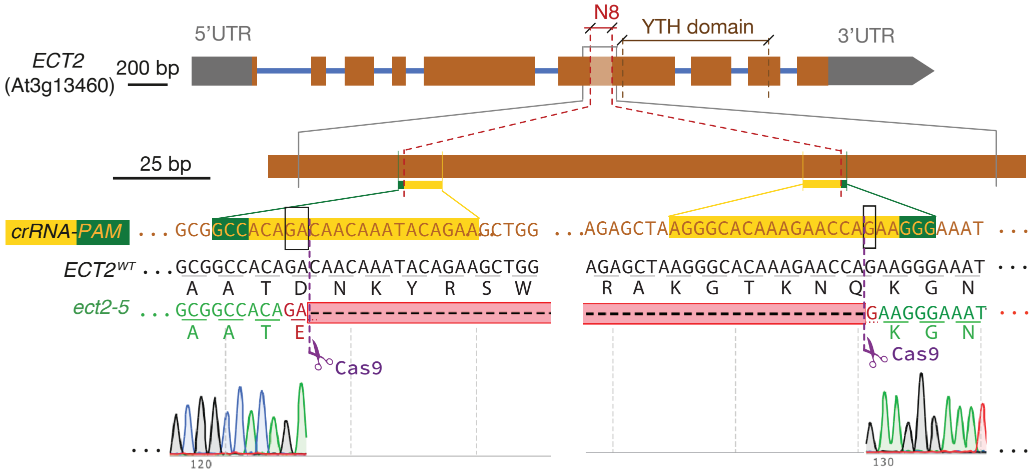


**Figure S1. CRISPR-Cas9 engineering of the *ect2-5* deletion mutant.**
Schematic representation of guide RNA design and the resulting in-frame chromosomal deletion matching nearly exactly the N8 element defined in transgenic deletion analysis[21]. crRNA, CRISPR-RNA, the sequence specificity components of the single-guide RNAs used to induce chromosomal *ECT2* deletions; PAM, protospacer adjacent motif.

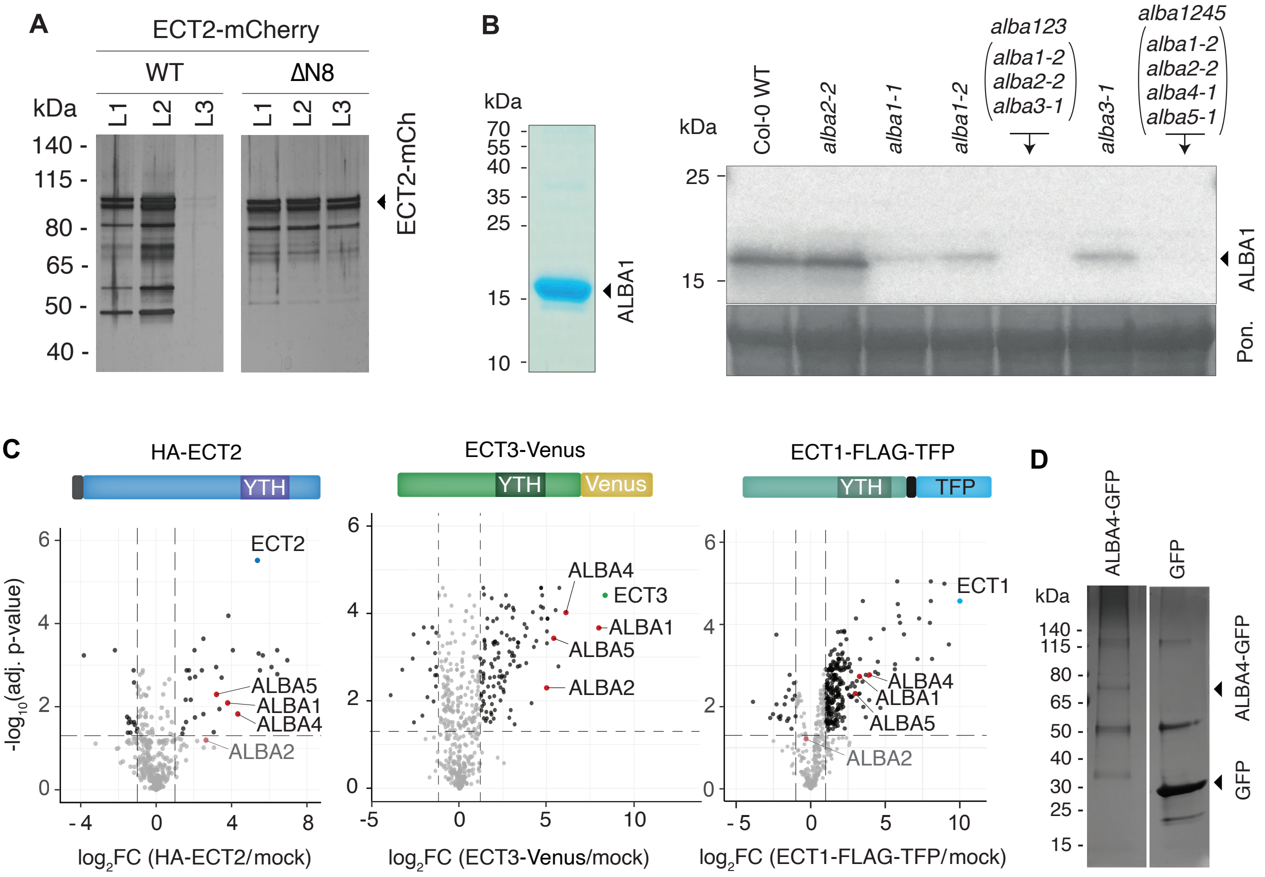


**Figure S2. Characterization of ECT2-ALBA interaction (supporting data).**

**(A)** Silver staining of aliquots of immunopurified fractions used for LS-MS/MS analysis of differential protein enrichment in ECT2-mCherry vs. ECT2$^{\Delta N8}$-mCherry purifications (supports **Figure 1G**).

**(B)** Left, coomassie stain of purified recombinant ALBA1 protein used for immunization of rabbits to produce antibodies. Right, western blot probed with the ALBA1 antibody. Ponceau staining is used as a loading control (supports **Figure 1H**). Note that although the *alba2-2* T-DNA allele used is not a full knockout, no ALBA2 protein is detectable in the *alba1245* mutant (see **Figure S5**).

**(C)** Volcano plot showing differential abundance of proteins immunopurified from total lysates of seedlings of the indicated transgenic lines compared to non-transgenic controls, highlighting the ALBA proteins identified in each case. Left, HA-ECT2[7] was purified with anti-HA beads; center, ECT3-Venus[7] was purified with GFP-trap; right, ECT1-TFP[20] was purified with GFP-trap. In all cases, proteins were identified and quantified by LC-MS/MS, and statistical significance was determined using empirical Bayes statistics with Benjamini–Hochberg adjusted p-values. These data have been published previously with no specific mention of ALBA proteins[21].

**(D)** Silver staining of aliquots of immunopurified fractions used for LS-MS/MS analysis of differential enrichment in ALBA4-GFP and GFP purifications (supports **Figure 1I**).

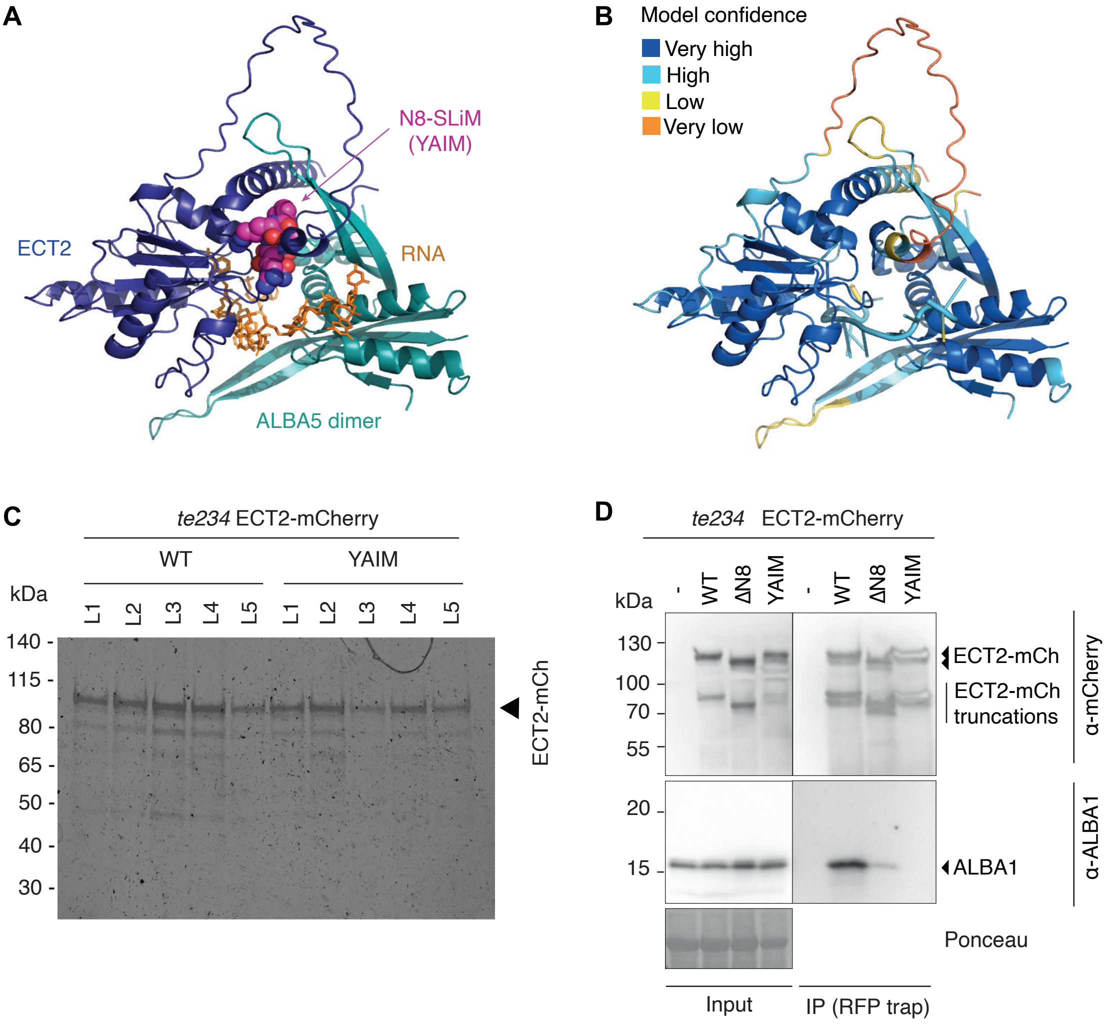


**Figure S3. The YTH-ALBA Interacting Motif (YAIM) is central for ECT2-ALBA interaction (supporting data).**
**(A)** Alternative view of the AlphaFold3 model shown in **Figure 2B** of the complex between ECT2 (YTH domain plus a YAIM-containing fragment of the N-terminal IDR), two ALBA5 subunits (ALBA domains only), and the 10-nt RNA [5'-AAA(m$^6$A)CUUCUG-3']. The YAIM is accentuated in space fill mode (Magenta, C; Blue, N; Red, O), all other protein elements in cartoon mode, and the RNA in stick mode (supports **Figure 2B**).
**(B)** Same view of the model as in panel (A), but colored according to the predicted Local Distance Difference Test (pLDDT) score calculated by Alphafold3 to indicate model confidence on a local per-residue basis[45] (supports **Figure 2C**).
**(C)** Silver staining of aliquots of immunopurified fractions used for LS-MS/MS analysis of differential enrichment in ECT2-mCherry and ECT2$^{YAIM}$-mCherry purifications (supports **Figure 2I**).
**(D)** Co-immunoprecipitation analysis of the ECT2-ALBA1 interaction. 9-day-old seedlings from three independent transgenic lines expressing each ECT2-mCherry variant were pooled prior to mCherry immunoprecipitation and analysis by western blot.

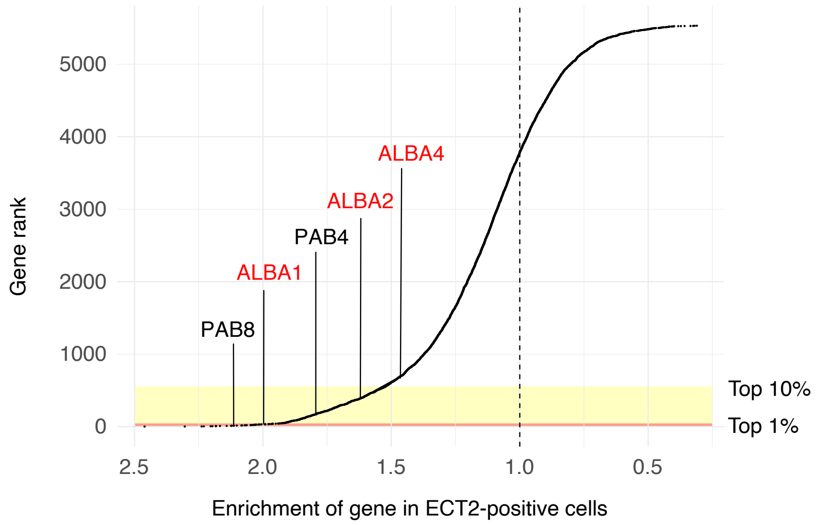


**Figure S4. Co-expression analysis of ECT2 and ALBA proteins.**
Odds ratio represents correspondence between ECT2-positive cells and cells expressing a given gene. Analysis controls for differences in UMI counts between ECT2-positive and ECT2-negative cells, and considers only genes expressed in between 20 and 80% of cells. Top 1% and 10% represents top genes out of those tested.

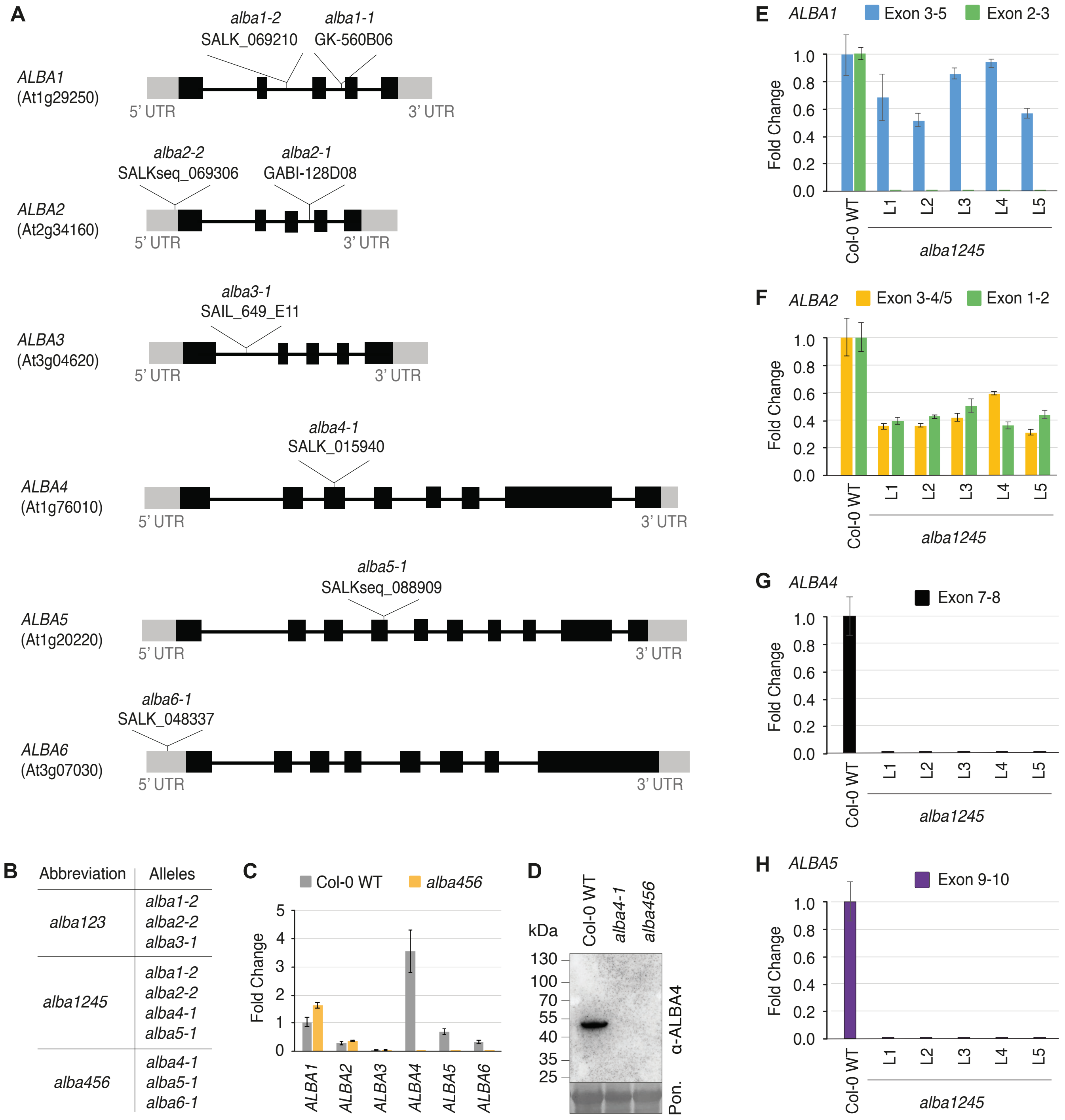


**Figure S5. Characterization of *ALBA* mutants (supporting data).**
**(A)** Schematic representation of *ALBA1-ALBA6* loci with the sites of T-DNA insertions indicated.
**(B)** Abbreviations of higher order *alba* mutants.
**(C)** *ALBA* mRNA levels measured by qPCR in Col-0 WT and *alba456*. Each measurement represents three biological replicates, with each replicate being composed of three individual plants. RNA levels were normalized to *CYCLOPHILIN* (At2g29960). Error bars represent the standard deviation of the means.
**(D)** Western blot probed with an ALBA4 antibody raised against synthetic ALBA4 peptides (see Methods). Ponceau (Pon.) staining is used as a loading control.
**(E-H)** *ALBA1, ALBA2, ALBA4* and *ALBA5* mRNA levels measured by qPCR in Col-0 WT and five individual *alba1245* plants, using primers spanning the indicated exons. RNA levels were normalized to *ACTIN2* (At3g18780). Error bars represent the standard deviation of the means.

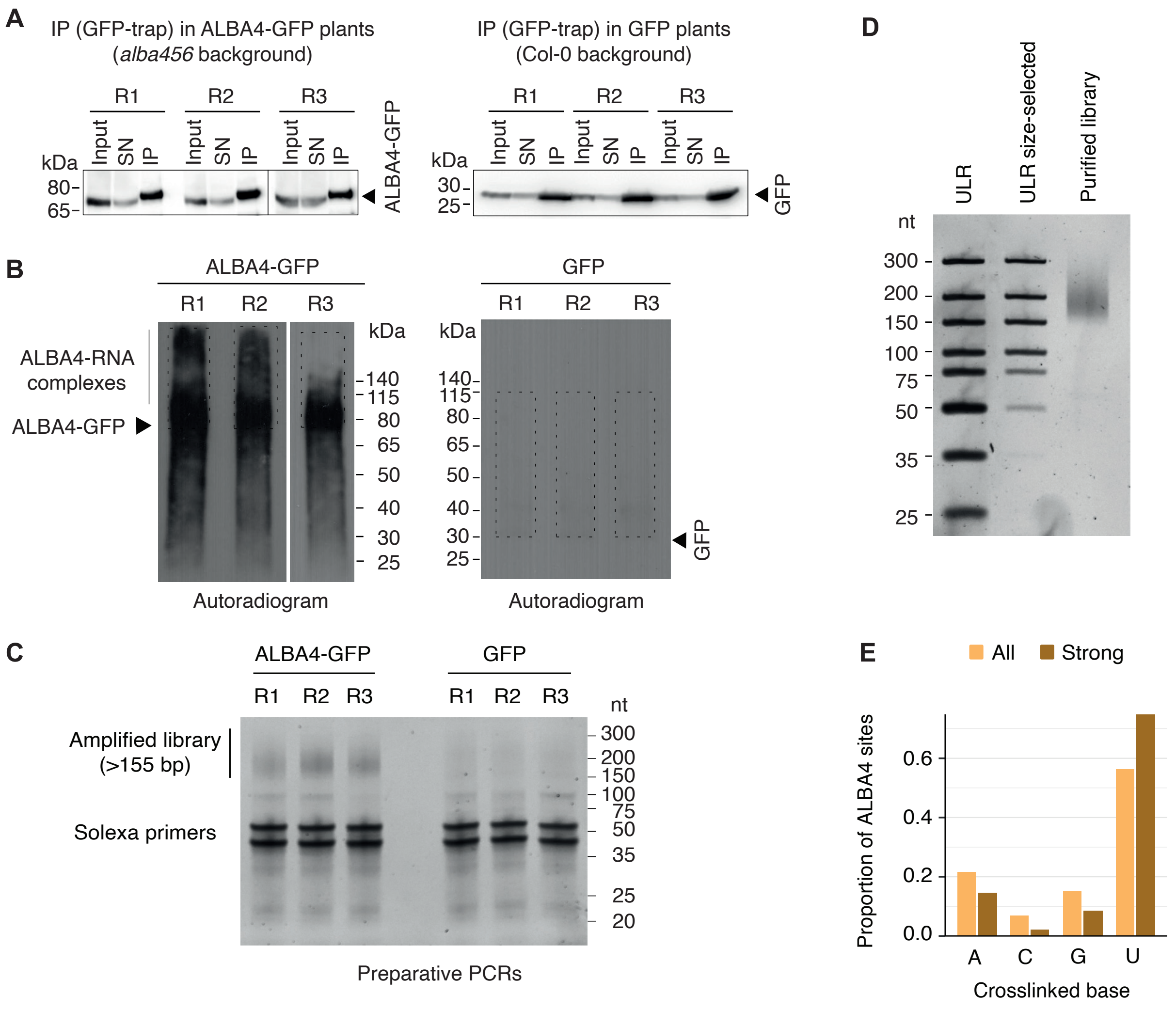


**Figure S6. ALBA4 target identification using iCLIP2 (supporting data).**

**(A)** Western blots against GFP after UV crosslinking and immunoprecipitation of RNA-protein complexes with GFP-trap beads in ALBA4-GFP plants (in *alba456* background) and GFP plants (in Col-0 background) (three replicates each). Presence of the respective protein is shown in the input (IN), supernatant (SN) after precipitation, and IP fraction.

**(B)** Autoradiogram of RNA-protein complexes from ALBA4-GFP (left) and GFP plants (right) after UV crosslinking and immunoprecipitation with GFP-Trap beads (three replicates each). Marker positions and the location of the ALBA4-GFP-RNA adducts are indicated. Dashed rectangles indicate the regions that were excised from the membrane. Autoradiograms of ALBA4-GFP were developed after 4 h exposure, while those of GFP were developed after overnight exposure.

**(C)** Gel electrophoresis of PCR-amplified iCLIP2 cDNA libraries (3 replicates each) visualized on a 12% polyacrylamide gel. A size standard indicates the fragment sizes. Libraries are visible as a smear at 155 nt and above.

**(D)** Gel electrophoresis of purified iCLIP2 cDNA library visualized on a 12% polyacrylamide gel. The cDNA library, alongside with the ultra-low range (ULR) molecular weight ladder, is subjected to size selection with ProNex beads to remove solexa primers. A ratio of ~15:1 between the 150-nt band and the 75-nt band of the URL ladder indicates efficient purification.

**(E)** Proportion of ALBA4 iCLIP2 crosslink sites according to the reference nucleotide, for both full and strong sets.

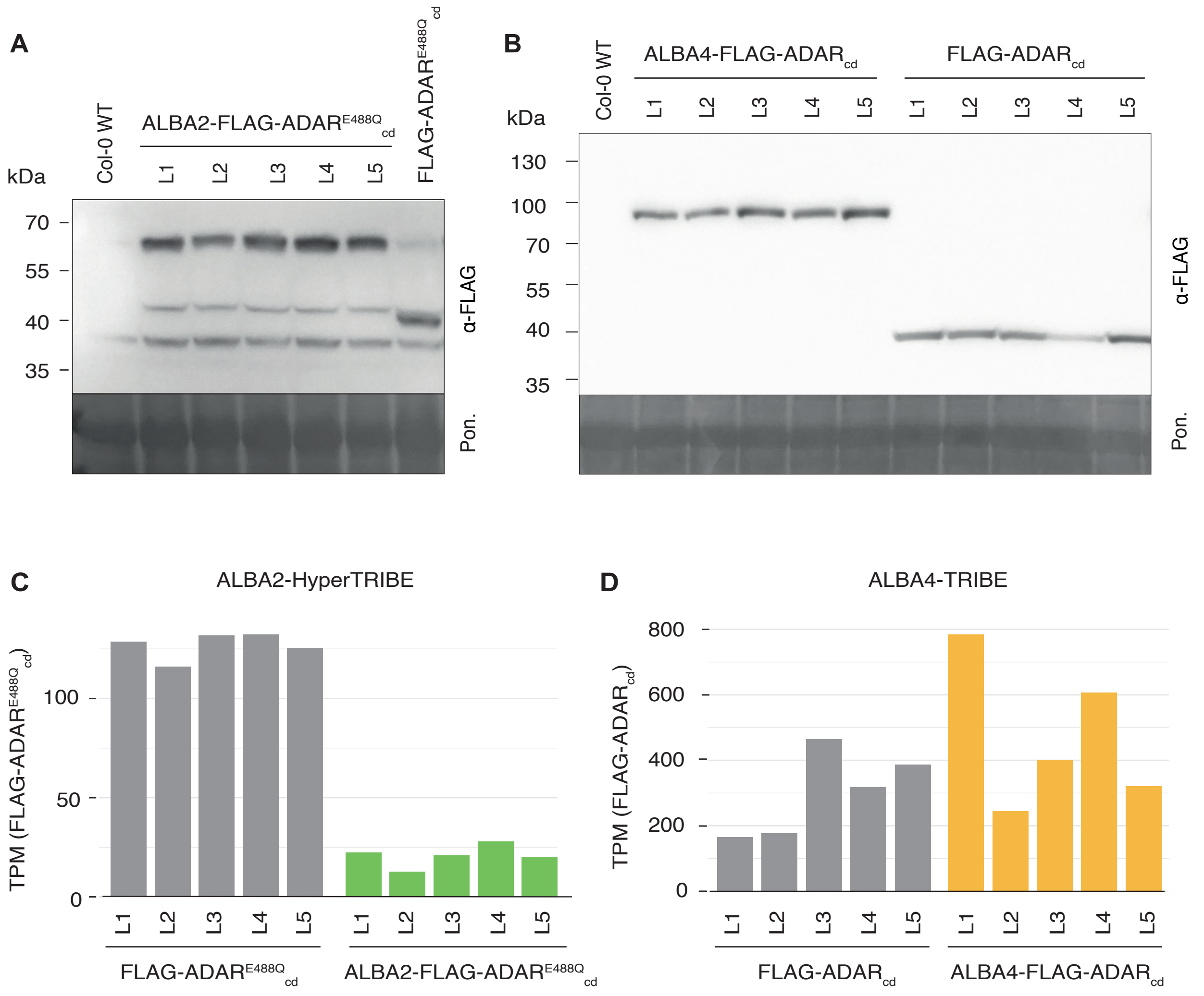


**Figure S7. ALBA2 and ALBA4 target identification by TRIBE and HyperTRIBE (supporting data).**
**(A)** Western blot of the independent lines selected for HyperTRIBE analysis of ALBA2. Ponceau (Pon.) staining is used as a loading control.
**(B)** Western blot analysis of the independent lines selected for TRIBE analysis of ALBA4. Ponceau (Pon.) staining is used as a loading control.
**(C)** Transcripts per million of FLAG-ADAR$^{E488Q}_{cd}$ detected in lines used for HyperTRIBE (ALBA2-HT).
**(D)** Transcripts per million of FLAG-ADAR$_{cd}$ detected in lines used for TRIBE (ALBA4-TRIBE).

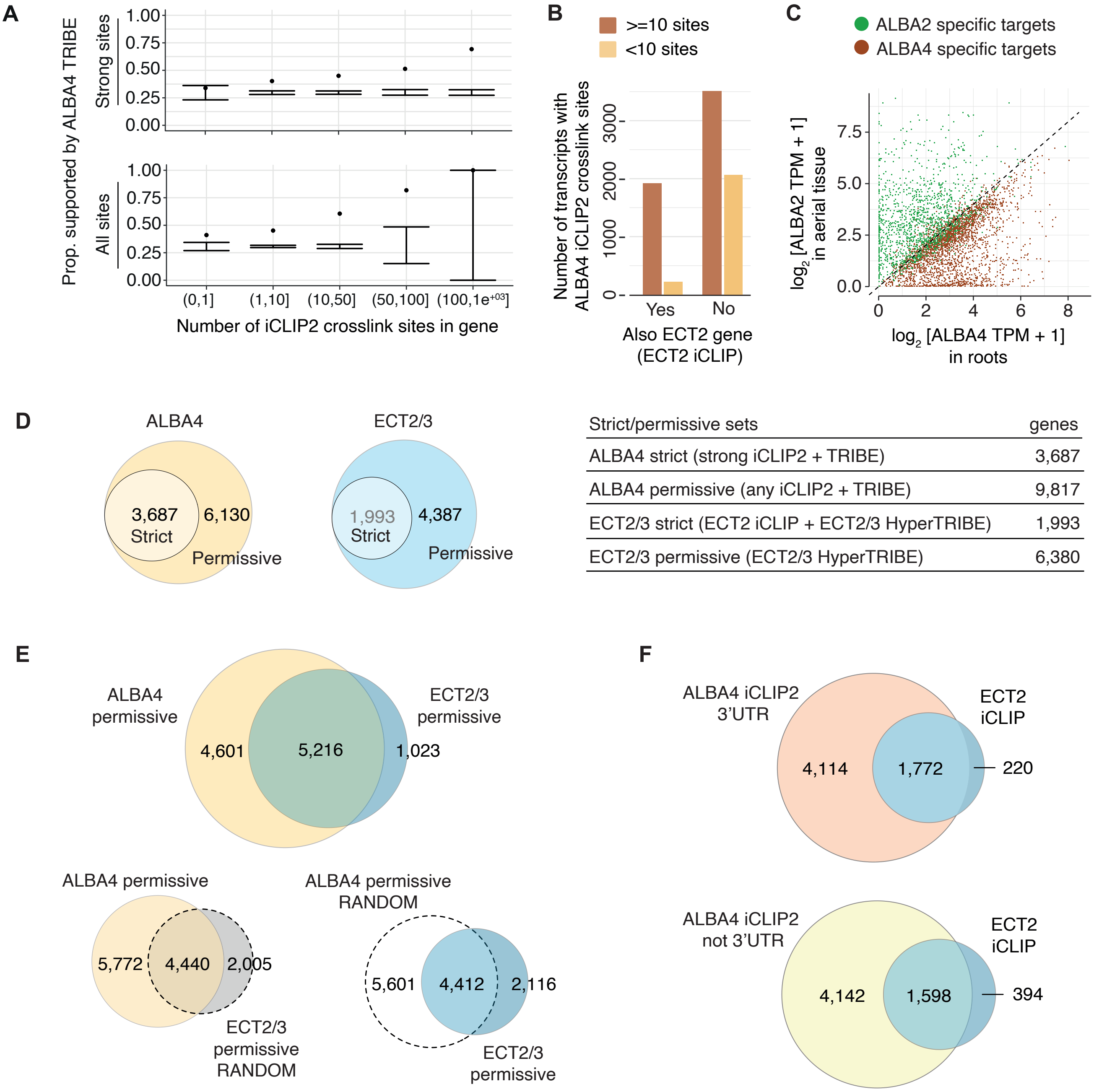


| Strict/permissive sets | genes |
|---|---|
| ALBA4 strict (strong iCLIP2 + TRIBE) | 3,687 |
| ALBA4 permissive (any iCLIP2 + TRIBE) | 9,817 |
| ECT2/3 strict (ECT2 iCLIP + ECT2/3 HyperTRIBE) | 1,993 |
| ECT2/3 permissive (ECT2/3 HyperTRIBE) | 6,380 |

**Figure S8. Analysis of ALBA2 and ALBA4 target sets (supporting data).**
**(A)** Proportion of genes supported by ALBA4-TRIBE according to the number of iCLIP2 crosslink sites. Points represent true proportions and intervals represent background distributed based on sampling genes of similar expression levels to the target genes from ALBA4 iCLIP2. Top panel represents the strong set of replicated ALBA4 iCLIP2 peaks and bottom panel represents the full set of replicated ALBA4 iCLIP2 peaks.
**(B)** Support of ALBA4 iCLIP2 site-containing genes according to whether the gene is also supported by ECT2 iCLIP.
**(C)** Scatter plot comparing expression for targets specific to either ALBA4 TRIBE (roots) or ALBA2 HyperTRIBE (aerial tissues). Values represent the average $log_2$(TPM+1) values from the Col-0 WT lines in each experiment.
**(D)** Table showing defined strict and permissive gene sets for ECT2/3 and ALBA4. Venn diagrams provide visual representations of the ECT2/3 and ALBA4 strict and permissive gene sets.
**(E)** Venn diagram for overlap between ALBA4 permissive and ECT2/3 permissive genes. Smaller Venn diagrams indicate overlap if either the ALBA4 or ECT2/3 permissive target sets were a randomly selected set of genes with a similar expression distribution to the true sets.
**(F)** Venn diagrams for overlap between ECT2 iCLIP-derived target genes (strict) and ALBA4 iCLIP2 genes, where the ALBA4 set either contains or does not contain crosslink sites in its 3'UTR.

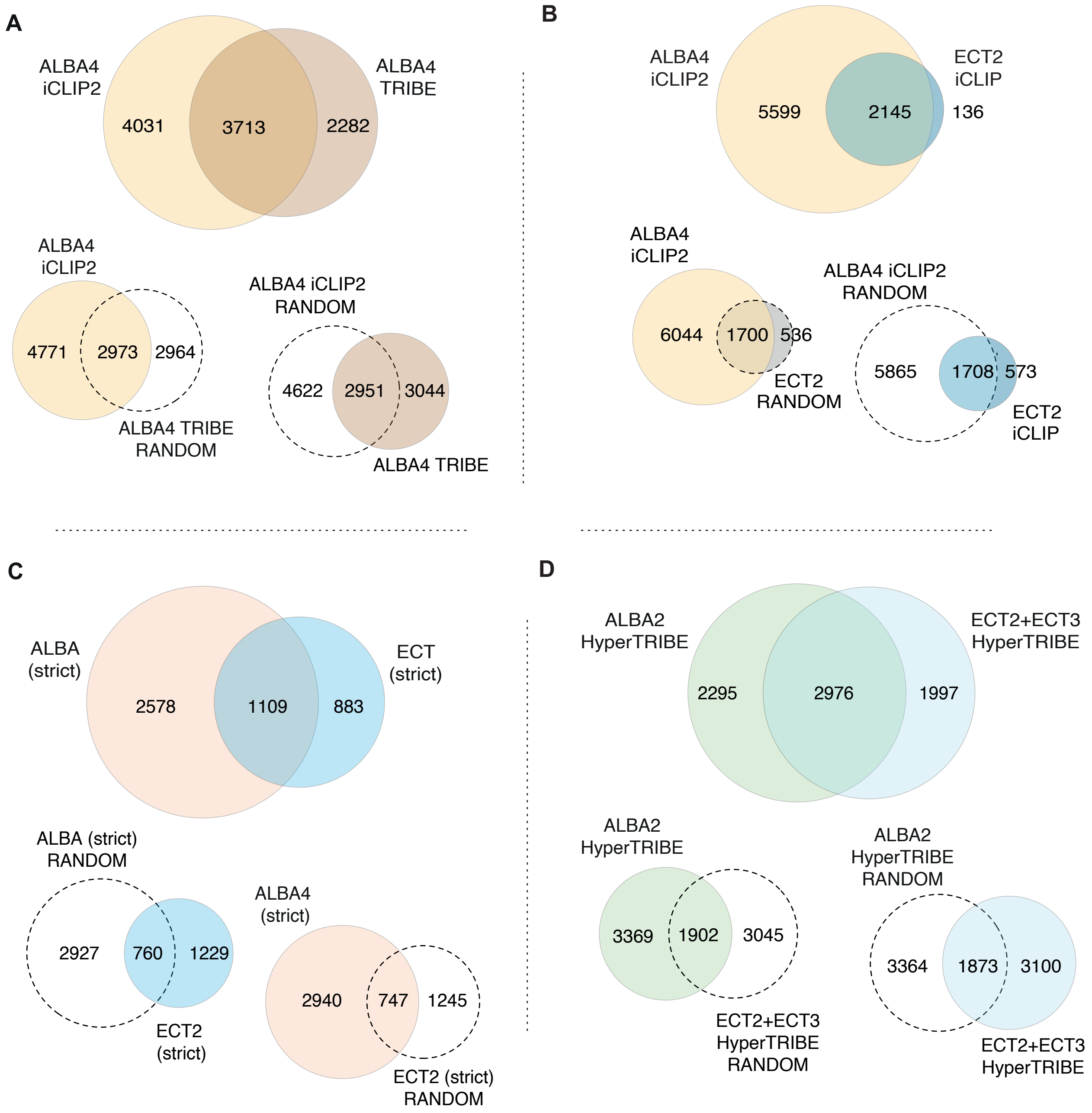


**Figure S9. Analyses of the overlap between ALBA2, ALBA4, ECT2 and ECT3 target sets determined by (Hyper)TRIBE and/or iCLIP.** Venn diagrams showing the overlap between the indicated target sets. In all cases, smaller Venn diagrams with one dashed circle indicate the overlap obtained if one of the sets used in the comparison were a randomly selected group of genes with a similar expression distribution to the true target set.

**(A)** Overlap between iCLIP2-defined and TRIBE-defined ALBA4 target sets.
**(B)** Overlap between iCLIP2-defined ALBA4 and iCLIP-defined ECT2 target sets.
**(C)** Overlap between strict ALBA4 (iCLIP2 + TRIBE support) and strict ECT2 (iCLIP + HyperTRIBE support) target sets.
**(D)** Overlap between ALBA2 and ECT2/3 HyperTRIBE target sets (only aerial tissues).

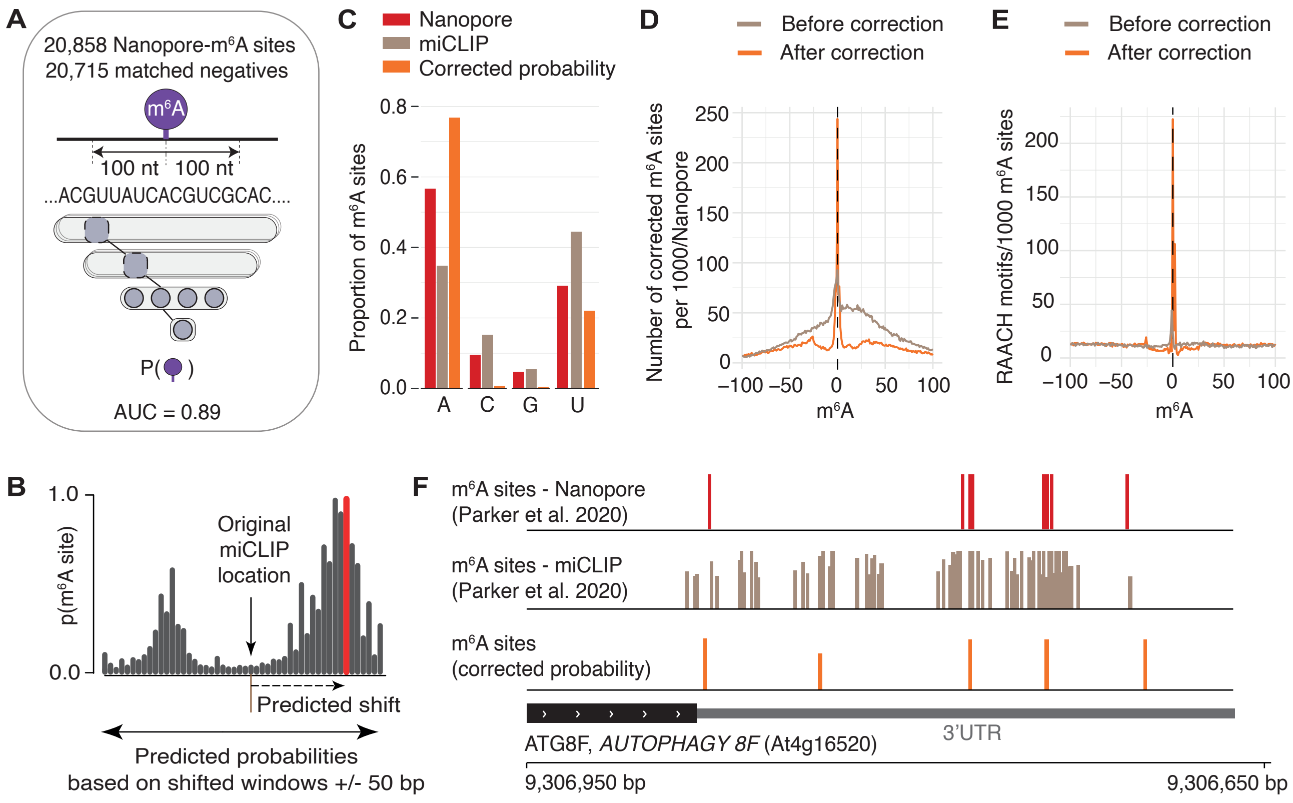


**Figure S10. A deep learning model to derive an augmented set of single-nucleotide-resolution m6A sites by integration of miCLIP and Nanopore data.**
**(A)** Strategy for the m6A deep learning model. Each Nanopore m6A site was paired with a location-matched negative, sequences +/- 100 bp from the site were extracted and used as input to a convolutional neural network tasked with predicting the presence or absence of m6A in the center of the input sequence. AUC = area under the curve.
**(B)** Strategy for correcting the exact position of m6A sites based on the deep learning model. For each site and all individual positions within a +/- 50 bp window, a +/- 100 bp sequence was extracted and used as input to the m6A-network to predict the presence of m6A. The position with the strongest prediction was chosen as the corrected position for that site.
**(C)** Reference base distribution of m6A sites as determined by Nanopore, miCLIP, or the corrected positions.
**(D)** Enrichment of Nanopore-determined m6A sites around the augmented set of m6A positions, before and after correction.
**(E)** Enrichment of RRACH motifs per 1000 augmented m6A sites, before and after correction.
**(F)** IGV view of a representative transcript, *AUTOPHAGY 8F* (At4g16520), showing the positions of m6A sites experimentally determined by either Nanopore or miCLIP, and the positions resulting from the correction.

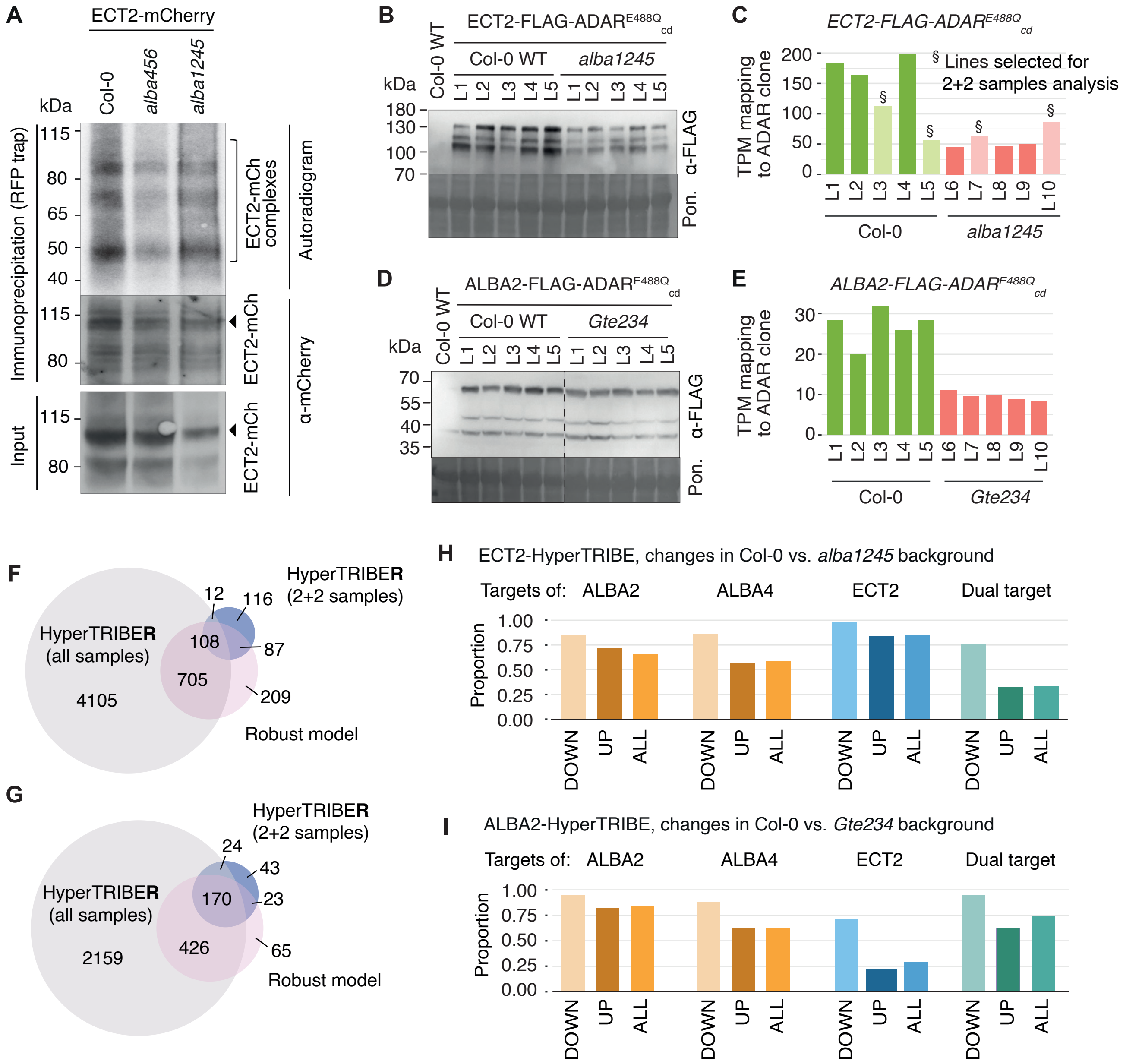


**Figure S11. Mutual ECT-ALBA dependence for target mRNA binding.**

**(A)** Autoradiogram (top) of RNA-protein complexes from ECT2-mCherry expressed in Col-0, *alba1245* or *alba456* after UV crosslinking and immunoprecipitation with RFP trap beads. Marker positions and the location of the ECT2-mCherry-RNA adducts are indicated. Immunoblots against mCherry show the precipitated protein in the IP (middle) and input (bottom). Samples were pools of 3 independent lines for each genotype.

**(B)** Western blot of the independent lines selected for HyperTRIBE analysis of ECT2 in Col-0 and *alba1245*. Ponceau staining was used as a loading control.

**(C)** Detected number of transcripts per million of ECT2-FLAG-ADAR in lines used for HyperTRIBE. Light bars indicate lines selected for the 2-sample analysis.

**(D)** Western blot of the independent lines selected for HyperTRIBE analysis of ALBA2 in Col-0 and *gte234*. Ponceau staining was used as a loading control.

**(E)** Detected number of transcripts per million of ALBA2-FLAG-ADAR in lines used for HyperTRIBE.

**(F)** Overlap of individual sites detected as significant in ECT2-HT in Col-0 vs. *alba1245* according to three different approaches - HyperTRIBER pipeline (no correction), HyperTRIBER pipeline (using lines indicated in C) and developed a robust model.

**(G)** As in (F) but counts depict containing genes for significant sites.

**(H-I)** Proportion of sites significantly differentially edited by ECT2- (H) or ALBA2- (I) -FLAG-ADAR$^{E488Q}_{cd}$ fusions in Col-0 vs. the indicated mutant background (*alba1245* in H; *Gte234* in I), which are targets of ALBA2, ALBA4, ECT2 or both ALBA4 and ECT2 (dual-targets). UP/DOWN categories are defined according to whether the $\log_2$ fold-change was negative (lower in the mutant background, DOWN) or positive (higher in the mutant background, UP).

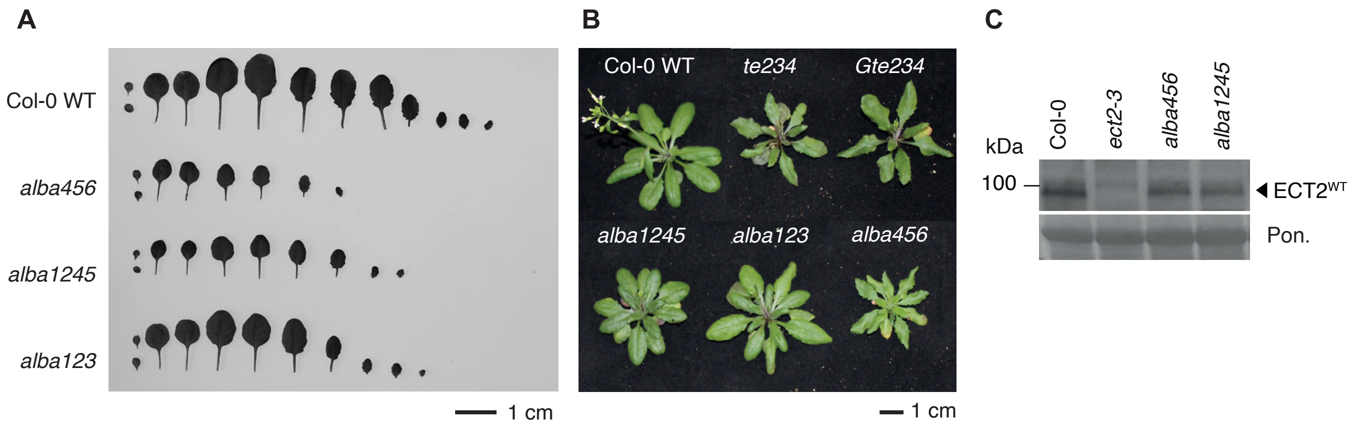


**Figure S12. Phenotypic analysis of *alba1245* and *alba456* mutants.**
**(A)** Leaf profiles of *alba* mutants at day 17 after germination.
**(B)** Delayed flowering phenotype of higher-order *alba* and *ect* mutants (5-week-old, germinated directly on soil).
**(C)** Protein blot of total lysates prepared from 10-day old seedlings of the indicated genotypes, probed with ECT2-specific antisera[7]. Arrow indicates the position of the ECT2$^{WT}$ protein. Ponceau staining serves as the loading control.
